# The trade-off function of photorespiration in a changing environment

**DOI:** 10.1101/2022.05.06.490933

**Authors:** Jakob Sebastian Hernandez, Thomas Nägele

## Abstract

The photorespiratory pathway in plants comprises metabolic reactions distributed across several cellular compartments. It emerges from the dual catalytic function of ribulose-1,5-bisphosphate carboxylase/oxygenase (Rubisco) which either carboxylates or oxygenates ribulose-1,5-bisphosphate (RuBP). Carboxylation reactions produce 3-phospho-glycerate (3PGA) molecules which are substrate for central carbohydrate metabolism while oxygenation forms 2-phosphoglycolate (2PG) molecules which are substrate for the multicompartmental recovery process of photorespiration. Further, 2PG is a strong inhibitor of several enzymes involved in the Calvin-Benson-Bassham cycle which challenges the experimental and theoretical study of carbon assimilation, photorespiration and metabolic regulation *in vivo*. Here, an approach of structural kinetic modeling (SKM) is presented to investigate the extend of stabilization of CBC and carbohydrate metabolism by photorespiration. Further, our approach highlights the importance of feedback regulation by 2-PG for alleviation of environmental perturbation. Our findings indicate that oxygenation of RuBP by Rubisco significantly stabilizes CBC activity and, thus, carbohydrate metabolism. Based on our findings, we suggest a trade-off function of photorespiration which reduces carbon assimilation rates but simultaneously stabilizes metabolism by increasing plasticity of metabolic regulation within the chloroplast. Furthermore, our analysis suggests a stabilizing effect of increasing the partition of newly assimilated carbon going towards sucrose biosynthesis. With this, our analysis sheds light on the role of a multicompartmental metabolic pathway in stabilizing plant metabolism within a changing environment.

## Introduction

Cellular metabolism consists of a highly elaborate reaction network which frequently shows complex and non-intuitive dynamical behavior. This behavior can be deterministically described by a system of ordinary differential equations, ODEs (Klipp et al., 2016). For example, such a kinetic model can then be used to analyze and simulate consequences of environmental fluctuations. After perturbations from its a steady state, a system can either return to its original state (stable), exponentially diverge from its original state (unstable), or be metastable (behavior is indifferent). The formulation of ODEs in a kinetic model is, however, typically challenged by lacking experimental data on enzymatic parameters, which are often laborious and difficult to quantify (Wittig et al., 2014). Furthermore, enzymatic activity *in vivo* depends on numerous factors, such as temperature and pH, thus greatly increasing the permissive parameter space which exacerbates the physiological interpretation of *in vitro* data (Bisswanger, 2017).

Previously, the approach of Structural Kinetic Modeling (SKM) was developed which allows for quantitative evaluation of dynamic properties without referring to any explicit system of ODEs, and which is achieved by a parametric representation of the Jacobian matrix (Steuer et al., 2006). SKM has been confirmed to be a suitable approach for evaluation of the stability and robustness of metabolic states (Grimbs et al., 2007). It has been successfully used in a variety of studies to analyze metabolism and its regulation (Grimbs et al., 2007; Reznik and Segrè, 2010; Fürtauer and Nägele, 2016). However, evaluation of regulatory interactions between a metabolite and reaction of interest is complicated by the complex interplay found within the highly dynamic system of cellular metabolism. Depending on the regulatory stetting, a single regulatory interaction can be beneficial or detrimental for the system stability. Further, although a vast amount of data and information is available on regulatory interplay between genes, proteins metabolites, e.g., provided by genome-scale metabolic networks (Tang et al., 2021), many regulatory and metabolic interactions can be assumed to remain elusive or unknown. Affirmatively, even in long studied and well-known biochemical reaction networks, new reactions or regulatory interactions are still being discovered. For example, within the Calvin-Benson Cycle (CBC) inhibition of sedoheptulose-bisphosphatase (SBPase) by 2-phosphoglycolate (2-PG) has recently been described (Flügel et al., 2017).

Carbon fixation takes place in the stroma of chloroplasts and is catalyzed by the enzyme ribulose-1,5-bisphosphate carboxylase-oxygenase (Rubisco). This enzyme facilitates the carboxylation of ribulose-1,5-bisphosphate (RuBP) and subsequent cleavage between the C2 and C3 carbon forming two molecules of 3-phospho-D-glycerate (3-PGA) (Andersson, 2008). In succeeding steps, 3-PGA is used in the CBC to resynthesize RuBP and supply the cell with triose phosphates, which act as precursors for starch and sucrose biosynthesis (Bassham et al., 1954; Martin et al., 2000). As the CBC is directly connected to the rate of carbon fixation and is ultimately the source of all carbohydrates used by the plant, its regulation is vital for overall metabolic stability. It thus has an immediate influence on the plants ability to cope with environmental fluctuations.

Perturbed or affected metabolic regulation can have drastic effects on the rate of carbon assimilation. For example, accumulation of triose phosphates through a low starch and sucrose synthesis rate would result in low abundance of inorganic phosphate and thus limit RuBP regeneration through a lower rate of photophosphorylation (Sharkey, 1985). In contrast, if triose phosphate utilization is too high, the rate at which RuBP needs to be regenerated is jeopardized as the CBC becomes carbon starved (Kadereit et al., 2014).

The rate at which Rubisco fixes carbon depends on both the CO_2_ and O_2_ concentration at the active site of Rubisco (Warburg, 1920; Foyer et al., 2009). This is a result of the dual catalytic function of Rubisco, which not only carboxylates RuBP, but also reacts with oxygen to form one molecule 2-PG and one of 3-PGA (Bowes et al., 1971; Foyer et al., 2009). As 2-PG is a strong inhibitor of several essential enzymes involved in the CBC and chloroplast metabolism, its degradation is of utmost importance (Anderson, 1971; Kelly and Latzko, 1976; Flügel et al., 2017). Affected enzymes are triose-phosphate isomerase (TPI) which catalyzes the interconversion between glyceraldehyde 3-phosphate (GAP) and dihydroxyacetone phosphate (DAHP) (Anderson, 1971), SBPase witch dephosphorylates sedoheptulose-1,7-bisphosphat (SBP) to sedoheptulose-7-phosphat (Sed7P) (Flügel et al., 2017), and phosphofructokinase (PFK) which catalyzes steps in glycolysis, (Kelly and Latzko, 1976).

The multistep pathway responsible for interconversion of 2-PG is known as photorespiration and involves several subcellular compartments (Fig. 1), namely the chloroplast, peroxisome, and mitochondria (Foyer et al., 2009). In this process, two molecules of 2-PG are converted into one molecule of CO_2_ and one molecule of 3-PGA which can then be used in the CBC and allows for the recovery of 75% of carbon otherwise lost (Berry et al., 1978; Foyer et al., 2009). This process, however, comes at a substantial energetic cost to the cell: fixed carbon is again released and ATP and NAD(P)H are consumed (Foyer et al., 2009). Furthermore, ammonium (NH_4_^+^) is produced during photorespiration and needs to be detoxified, adding further costs to the cell in the form of ATP and reduced ferredoxin (Foyer et al., 2009).

**Fig. 1:**
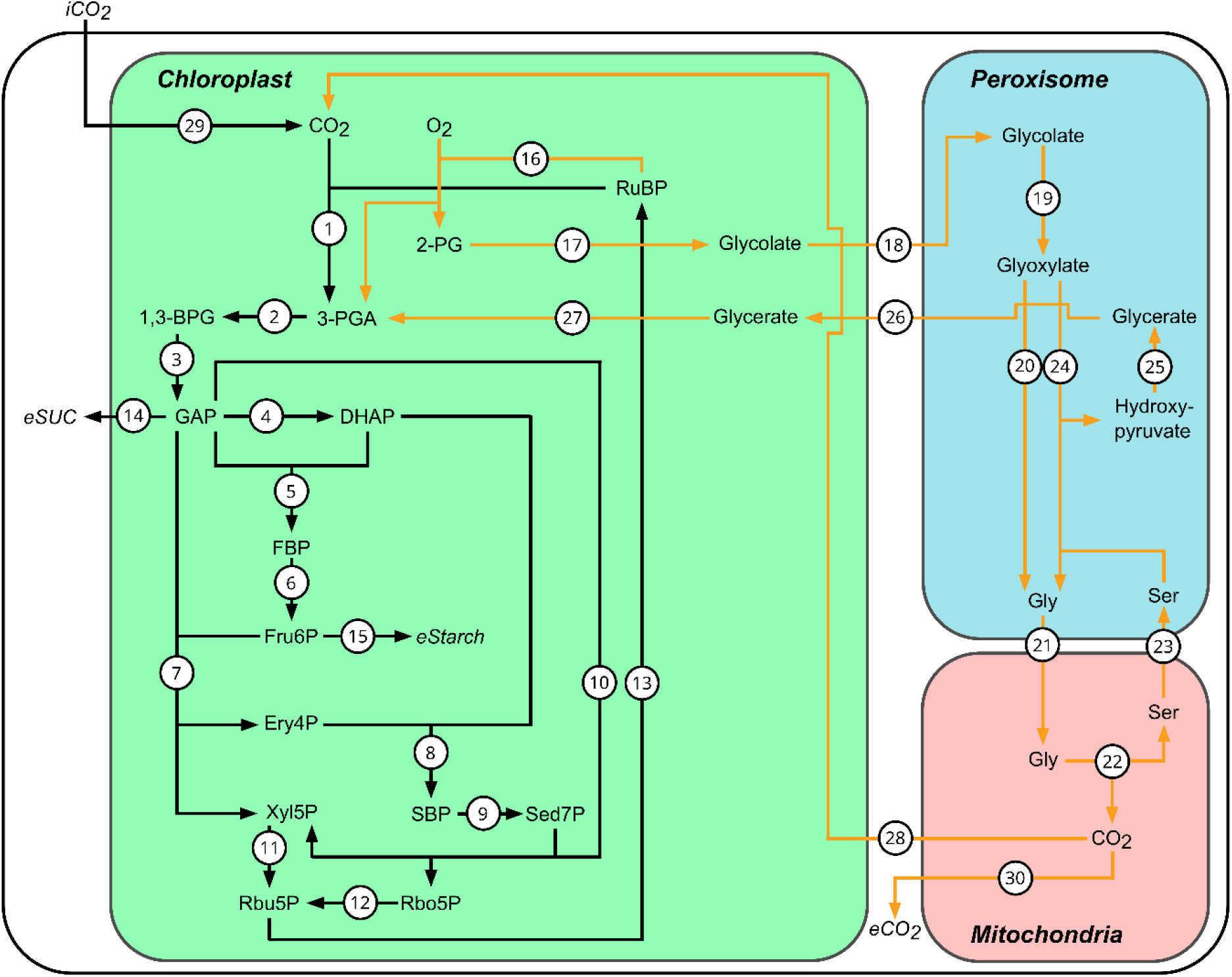
The Calvin-Benson Cycle (black) and photorespiration (orange). Reactions are catalyzed by the following enzymes: 1. Rubisco – ribulose-1,5-bisphosphate carboxylase-oxygenase, 2. PGK – Phosphoglycerate kinase, 3. GAPDH – Glyceraldehyde 3-phosphate dehydrogenase, 4. TPI – Triose-phosphate isomerase, 5. ALD – Aldolase, 6. FBPase – Fructose 1,6-bisphosphatase, 7. TK – Transketolase, 8. ALD – Aldolase, 9. SBPase – Sedoheptulose-1,7-bisphosphatase, 10. TK – Transketolase, 11. RPE – Ribulose-phosphate epimerase, 12. RPI – Ribose-5-phosphate isomerase, 13. PRK – Phosphoribulokinase, 14. GAP export for sucrose synthesis, 15. Fru6P export for starch synthesis, 16. Rubisco – ribulose-1,5-bisphosphate carboxylase-oxygenase, 17. PGP – Phosphoglycolate phosphatase, 18. Glycolate transport, 19. GO – Glycolate oxidase, 20. GGAT – Glutamate-glyoxylate aminotransferase, 21. Glycine transport, 22. GDC and SHMT – Glycine decarboxylase complex and Serine hydroxymethyltransferase, 23. Serine transport, 24. SGAT – Serine-glyoxylate aminotransferase, 25. HPR1 – Hydroxypyruvate reductase 1, 26. Glycerate transport, 27. GLYK – Glycerate kinase, 28. Portion of CO_2_ reentering chloroplast, 29. CO_2_ entering chloroplast form atmosphere, 30. Portion of CO_2_ released to cytoplasm. RuBP – Ribulose 1,5-bisphosphate, 3-PGA – 3-Phospho-D-glycerate, 1,3-BPG – 1,3-Bisphosphoglyceric acid, GAP – Glyceraldehyde 3-phosphate, DHAP – Dihydroxyacetone phosphate, FBP – Fructose 1,6-bisphosphate, Fru6P – Fructose 6-phosphate, Ery4P – Erythrose 4-phosphate, Xyl5P – Xylulose 5-phosphate, SBP – Sedoheptulose 1,7-bisphosphate, Sed7P – Sedoheptulose 7-phosphate, Rbu5P – Ribulose 5-phosphate, Rbo5P – Ribose 5-phosphate, 2-PG – 2-Phosphoglycolate, Gly – Glycine, Ser – Serine. The model does not consider refixation of ammonium.

Historically, oxygenation of RuBP was viewed as an unavoidable consequence of an oxygen rich atmosphere, with photorespiration only function being the disposal of 2-PG (Lorimer and Andrews, 1973). However, an increasing body of knowledge points to a more essential role of photorespiration besides its function in metabolic repair. To this extend, it has been suggested that photorespiration has a significant impact on protection from photoinhibition (Heber and Krause, 1980; Takahashi et al., 2007; Shi et al., 2022).

As regulation of the CBC is crucial for stability of whole plant metabolism, this study aimed to investigate and quantify to what extend photorespiration affects the stability of the CBC. Additionally, it was questioned if 2-PG, in its function as a key regulator, can alleviate environmental perturbations. For its analysis, a discreet-parameter-optimization approach is introduced beings based on a modified version of the Branch and Bound (BnB) algorithm (Land & Doig, 1960). To demonstrate this approach, we reevaluated the stability of CBC, using SKM to consider the effect of photorespiration.

## Materials and Methods

### Stability of reaction networks

Biochemical reaction networks can be deterministically described over time ***t*** by a system of ordinary differential equations (ODEs), 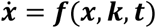 with ***x*** = (*x*_1_, …, *x*_*n*_)^*T*^ variables (e.g. substrate concentrations), ***f*** = (*f*_1_, …, *f*_*n*_)^*T*^ functions, and ***k*** = (*k*_1_, …, *k*_*l*_)^*T*^ parameters (e.g. kinetic constants) (Klipp et al., 2016).

If the point ***x*** fulfills the steady state condition 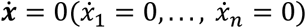, the system will remain in its current state without external perturbations and ***x*** is annotated as 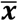. If, however, the system ***x*** is perturbated the steady state can either be stable (***x*** returns to steady state), unstable (***x*** leaves steady state), or metastable (behavior is indifferent) depending on the system behavior (Klipp et al., 2016).

The dynamics of an ODE system 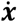 close to a stable point 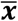 can be investigated by means of linearization. To this end, the Jacobian Matrix ***J*** is evaluated at 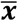 and can be obtained by Taylor expansion of temporal changes of the deviations from the steady state 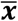. The best linear approximation of 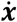 near 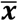 is then given by 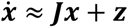 with ***z*** = (*z*_1_, …, *z*_*n*_)^*T*^ containing inhomogeneities (Klipp et al., 2016). For non-homogenous systems (i.e. ***z*** ≠ 0) the linearization can be converted into a homogenous system by the coordination transformation 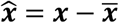 resulting in 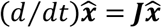 and the general solution for homogenous systems:

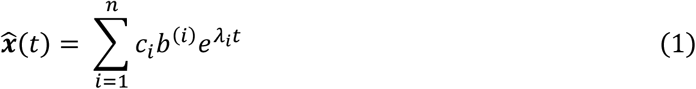

with *λ*_*i*_ being the eigenvalues of ***J*** and *b*^(*i*)^ being the corresponding eigenvectors (Klipp et al., 2016). The coefficients *c*_*i*_ can be determined by solving 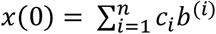 (Note that 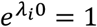). As a consequence of *e*^*λt*^ = *e*^(*a*+*ib*)*t*^ = *e*^*at*^(cos *bt* + *i* sin *bt*), the stability of the steady state is entirely determined by the real parts of the eigenvalues of ***J***. If all real parts of the eigenvalues are below zero (i.e. *a* < 0) then for *t* → ∞ the term *e*^*λt*^ converges to 0 and thus 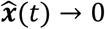and 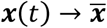. In other words, after a perturbation the system returns to 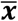 and the steady state is considered locally asymptotically stable. On the other hand, if any of the eigenvalues has a positive real part (i.e. *a* > 0) the system will exponentially diverge from its steady state.

An impotent note is that, following the Hartman–Grobman theorem, the linearized system only reflects the local behavior of the dynamic system if the stable point 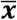 is hyperbolic (i.e. ***J*** cannot have eigenvalues whose real parts equal 0, *a* ≠ 0).

### Structural Kinetic Modeling

In practice, calculation of the Jacobian Matrix ***J*** is often complicated by a lack of knowledge on enzymatic rate equations and kinetic parameters (e.g. *V*_*max*_, *K*_*m*_) required for the formulation of the ODE system 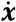. To circumvent this, an approach dubbed “structural kinetic modeling” (SKM) was proposed by Steuer and co-workers in which kinetic parameters are replaced by normalized enzymatic parameters (Steuer et al., 2006).

In this approach the system of ODEs is given by:

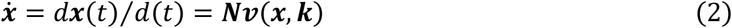

with ***x*** = (*x*_1_, …, *x*_*n*_)^*T*^ metabolites and ***v*** = (*v*_1_, …, *v*_*r*_)^*T*^ reaction rates. The *n* x *r* dimensional matrix ***N*** represents the stochiometric matrix of the considered metabolic reaction network. After defining:

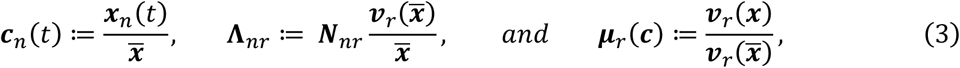

the variable substitution 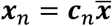 leads to the system being rewritten in terms of ***c***(***t***) as:

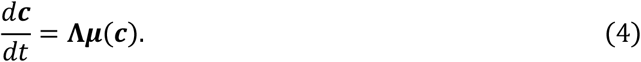

Now ***J*** can be evaluated at steady state ***c***^0^ = 1 with:

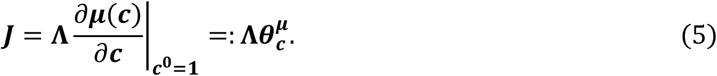

Thus, the Jacobian is defined as a product of matrices **Λ** and ***θ***. The matrix **Λ** consists of the stochiometric matrix ***N*** normalized to the flux 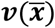 and metabolite concentrations 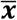 at steady state. The *r* x *n* dimensional matrix ***θ*** contains normalized elasticities and represents the degree of saturation of the normalized flux ***μ***(***c***) with respect to the normalized concentration ***c***(*t*). In other words, entries in ***θ*** describe to what extend changes in concentration of metabolite *n* influence the flux rate of reaction *r*. Depending on the interaction between the considered metabolite and reaction, ***θ*** is either defined within the interval (0; 1] for activating effects or within (−1; 0) for inhibitory effects. A detailed discussion concerning entries of ***θ*** is given in (Steuer et al., 2006). Entries in relation to Michaelis-Menten Kinetics are discussed in (Reznik and Segrè, 2010).

To analyze the system dynamics, the model is then repeatedly simulated with parameters for 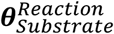 being chosen from a uniform distribution in the unit interval (0; 1]. This ensures that all possible explicit kinetic models of the observed system are considered. The Eigenvalues of each randomized Jacobian is calculated and the stability of the model is then evaluated.

### Ranking of parameters

Parameters used in the SKM approach are ranked according to their Pearson correlation to *max*(*Re*(*λ*)) and the distance between the distribution function of stable solution and the original probability density function. Pearson correlation is calculated as follows:

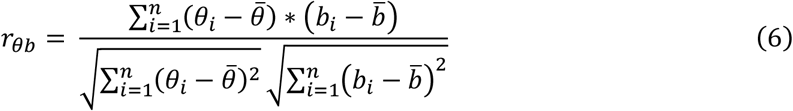

with *b* = *max*(*Re*(*λ*)). The rank for the correlation analysis *R*_*c*_ for a Parameter 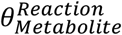 is given in descending order of |*r*_*θb*_|. The distance is calculated by:

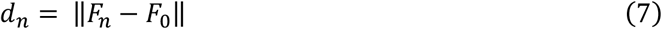

and ranked in descending order of *d*_*n*_. The overall rank for a parameter is given in ascending order of the mean from the ranks *R*_*c*_ and *R*_*d*_.

### Branch and Bound algorithm

To analyze how regulation affects the observed stability of a model, a discreate parameter optimization algorithm has been developed. It follows the *Branch and Bound* procedure first proposed by (Land and Doig, 1960), which allows the implicit enumeration of all possible combinations of regulatory interactions in a given model. The algorithm uses a best-first search strategy with wide branching as reviewed before (Morrison et al., 2016). An added challenge arises from the fact that the objective of the optimization problem (proportion of stable solutions) is not easily calculated by a simple cost function but rather has to be determined using a Monte Carlo experiment which is inherently noisy. Owing to this, nodes are pruned by hypothesis testing. As the proportion of stable solutions follows a binomial distribution which can be normally approximated, the Z-test was used (***H***_**0**_: *p*_1_ ≤ *p*_2_, the proportion of stable solutions of the heuristic solution is equal or less than that of the considered node. ***H***_**1**_: *p*_1_ > *p*_2_, the proportion of stable solutions of the heuristic solution is greater than that of the considered node). The standard score was calculated with:

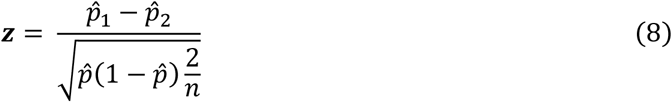

with 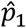 being the proportion of stable solutions within the best heuristic solution (i.e. the best solution with only discreet parameters). The proportion of stable solutions in the node considered for pruning is given by 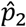. The pooled standard error was calculated using 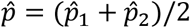. The number of samples within each Monte Carlo experiment is given by *n*. If the value of ***z*** > 2.33 (*α* = 0.01, one-tailed), then ***H***_**0**_ can be discarded (i.e. the proportion of stable solutions in the heuristic solution is significantly higher than in the observed node) and the considered node is pruned.

For continuous parameter optimization of nodes, particle swarm optimization was applied (Kennedy and Eberhart, 1995). The velocity 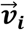 and position 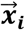 of a particle *i* is updated according to:

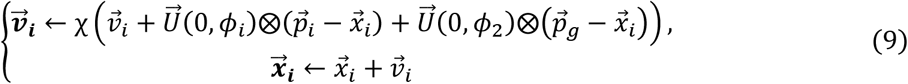

with *ϕ* = *ϕ*_1_ + *ϕ*_2_ = 4.1, *ϕ*_1_ = *ϕ*_2_ and 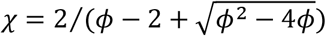 as proposed by (Clerc and Kennedy, 2002) and reviewed in (Poli et al., 2007).

Both pseudocode of the algorithm and an accompanying MATLAB® (www.themathworks.com) script are provided in the supplementary material.

## Results

### Photorespiration affects stability of carbon fixation

A simplified mathematical model was developed to simulate fluxes under the steady state assumption 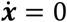. The considered equilibrium point 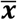 was hyperbolic and in accordance with the Hartman– Grobman theorem could be used to analyze the local behavior of the dynamical system (Hartman, 1960), justifying the use of the SKM approach.

Based on real parts of eigenvalues of Jacobian matrices, the stability of the model was then analyzed and compared to a model in which Rubisco only catalyzes the carboxylation of RuBP (no photorespiration). In a synthetic scenario, stability was analyzed under conditions where 2-PG had an unspecific efflux and no carbon was recovered. A graphical representation of these models can be found in the supplementary material (Fig. S1). Model simulations were based on literature data and run 10^6^ times (Zhu et al., 2007; Jablonsky et al., 2011; Flügel et al., 2017; Fürtauer et al., 2019; Küstner et al., 2019; Kitashova et al., 2021).

In the most basic case of all 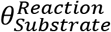 being set to 1 (corresponding to simple irreversible mass-action kinetics) the full model (including all reactions shown in Figure 1) and the model with an unspecific 2-PG efflux showed a stable steady state. Strikingly, the model without photorespiration demonstrated an unstable steady state (Fig. 2).

**Fig. 2:**
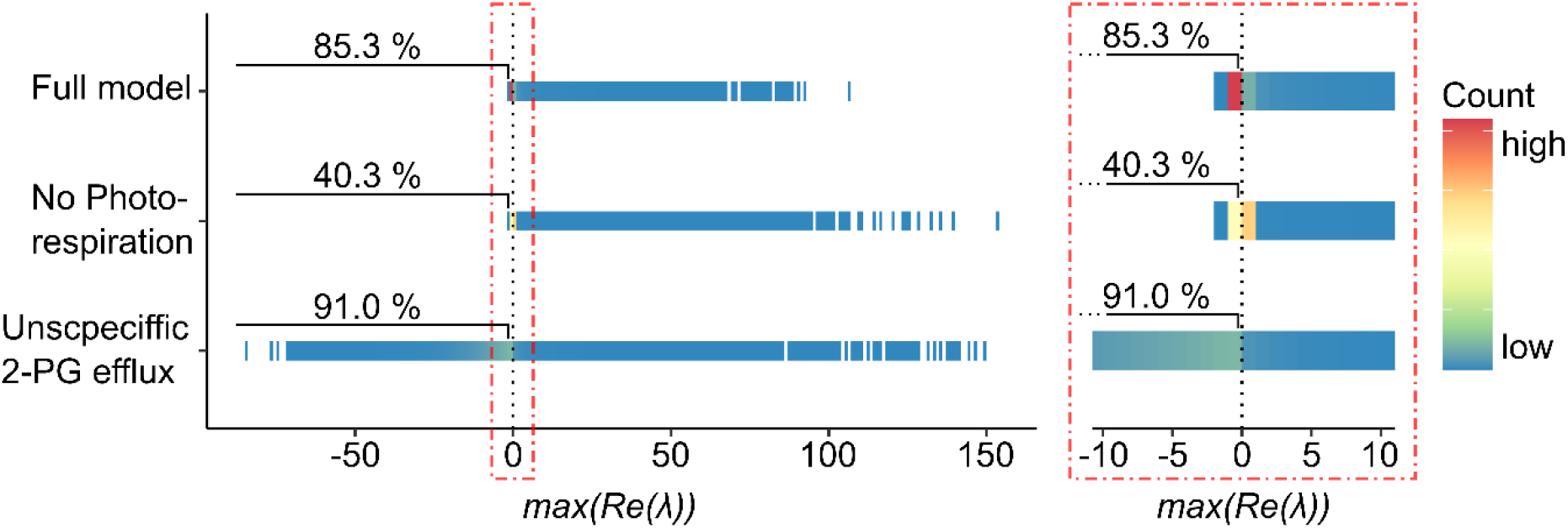
Stability of the CBC. Results are shown for models with photorespiration, without photorespiration, and with an unspecific 2-PG efflux. Maximal real eigenvalues below 0 (dotted black line) correspond to stable systems. The range of maximal real parts of eigenvalues between -10 and 10 is enlarged and is shown on the right. Each model was simulated 10^6^ times. Input and saturation parameters where randomized.

When allowing 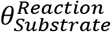 to move freely in the interval (0; 1], the results indicated that the model without photorespiration was the least stable with only 40.3 % of simulations resulting in a stable solution, i.e., *max*(*Re*(*λ*)) was negative (Fig. 2). Comparison to the full model showed that including the photorespiratory pathway, the system stabilized to 85.3 %. Interestingly, the most stable model was observed with an unspecific efflux of 2-PG, in with 91.0 % of maximal real parts of eigenvalues where blow 0. This suggested that oxygenation of RuBP itself increases stability of the CBC.

### GAP dynamics are pivotal for system stability

In order to investigate where the 14.7% instability observed in the full model originated from, the perturbated parameters of the model where ranked in accordance to their impact on the stability of the metabolic network. To this extend, the Pearson correlation between parameters and *max*(*Re*(*λ*)) was determined. Furthermore, the distribution of perturbed parameter values resulting in a stable solution was compared to their original probability density function, *d*_*n*_ = ‖*F*_*n*_ − *F*_0_‖. Of the most influential parameters, almost all were found to be tied to reactions and metabolites within the CBC, except for 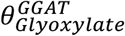 and 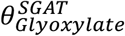 which were associated with photorespiration (Fig. 3A). Based on these findings, the most important reactions where identified (red marks in Fig. 3B). The results showed that reactions involving GAP as a substrate as well as the carboxylation and oxygenation of RuBP were of particular importance for stabilization of system behavior after perturbation. In conclusion, this suggests GAP partitioning to be essential for CBC stability.

**Fig. 3:**
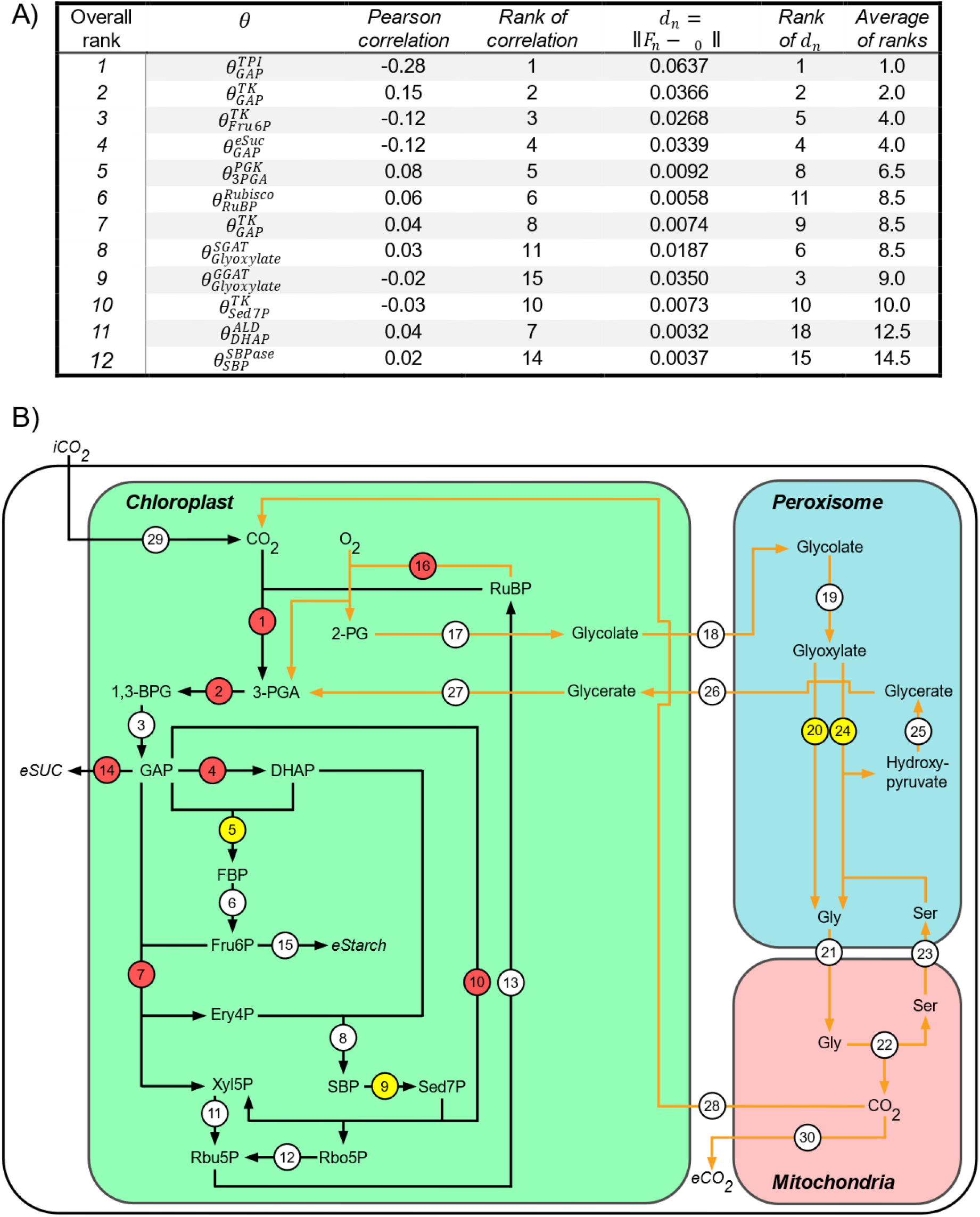
Parameters influencing the stability of the full model. A) Ranking of the 12 most influential parameters according to their correlation to *max*(*Re*(*λ*)) and distance *d*_*n*_. B) Visualization of the most important reactions according to their impact on the stability of the model. The most influential reactions are indicated in red. Reactions of medium importance are indicated in yellow. Reactions of little importance to stability are indicated in white.

Interestingly, reactions catalyzed by TPI, and to a lesser extend also SBPase, had a considerable impact on the stability of the model. This further suggested a central regulatory role of 2-PG which inhibits both enzymes.

The same analysis was repeated for the model lacking photorespiration (Fig. S2). Strikingly, there was a sharp increase in the value of *d*_*n*_ for two parameters, 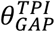 and 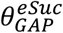. The *d*_*n*_ value for 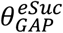 increased from 0.037 to 0.289, for 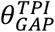 the change was slightly lower, increasing from 0.067 to 0.245. As *d*_*n*_ represents a shift in the parameter distribution found in stable solutions from the uniform distribution, the values of *θ* contained in stable solutions where compared between the two models (Fig. 4).

**Fig. 4:**
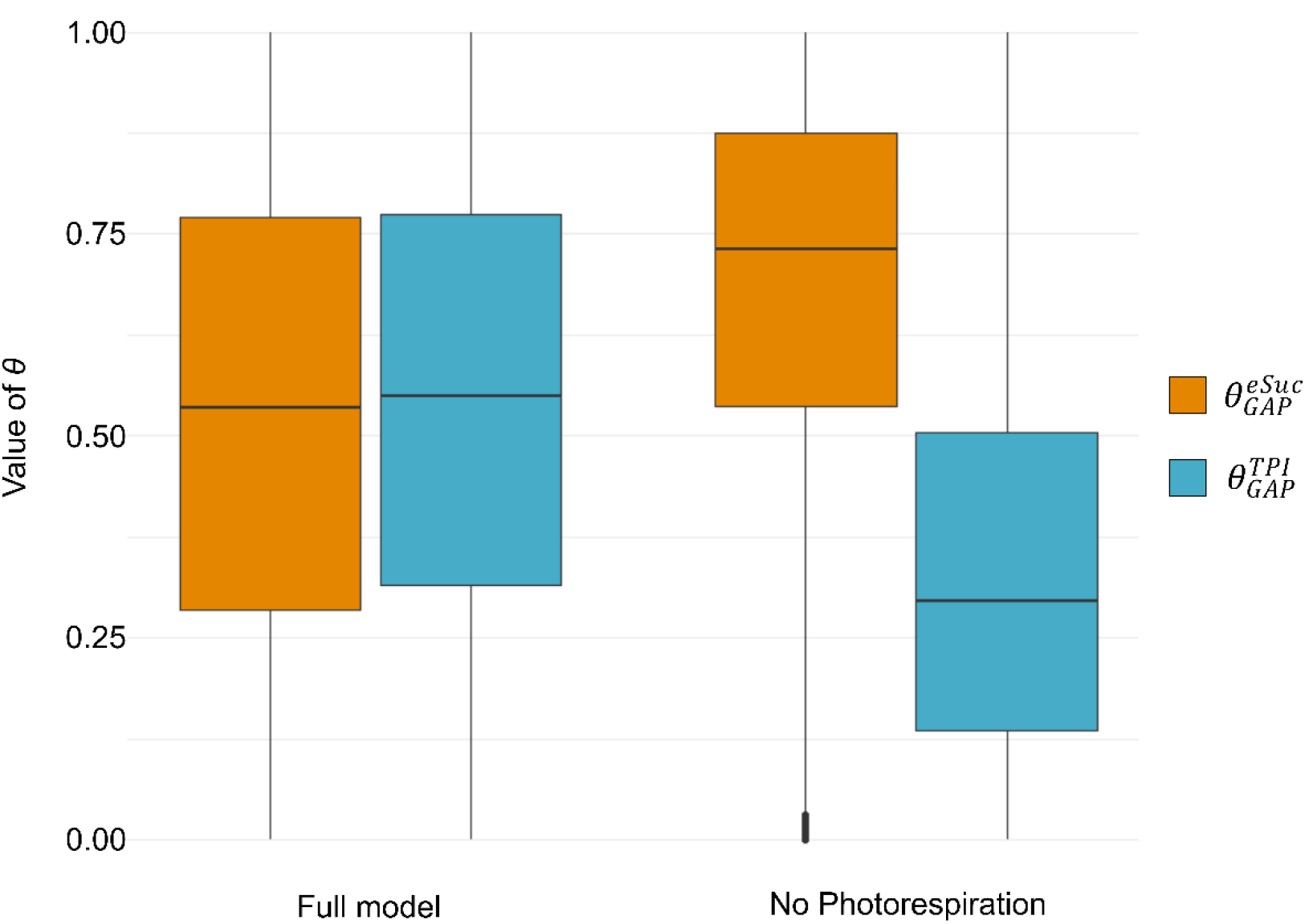
Values of *θ* found in stable solutions. Parameters are shown in which the model without photorespiration showed the largest deviation from the expected distribution 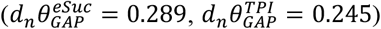. Values for *θ* are shown for both the full model and the model excluding photorespiration.

The results showed that a lack of photorespiration clearly reduces the parameter space of stable solutions compared to the full model. Medians of the saturation parameter 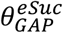 were shifted towards 1, while 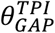 was shifted towards 0. This suggested an increasing importance of GAP export in order to stabilize the system, especially in relation to GAP utilization for DAHP synthesis. As starch synthesis occurs downstream from TPI, this indicated that starch synthesis may not be able to stabilize fluctuations effectively. Taken together, the structure and location of photorespiration thus appears to alleviate the pressure on the GAP export required to stabilize the system.

To investigate if starch synthesis is indeed unable to affect the stability of the system, the effect of 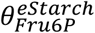 on the stability was analyzed across all models (Table I).

**Table I:** Effect of 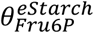 on model stability.

| *Model* | *Rank* | *Pearson Correlation* | *d<sub>n</sub>* |
| --- | --- | --- | --- |
| *Full model* | 32 of 34 | 0.00 | 0.0018 |
| *No Photorespiration* | 12 of 20 | 0.00 | 0.004 |
| *Unspecific 2-PG efflux* | 19 of 22 | -0.01 | 0.001 |

In all three models, 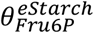 showed no apparent effect on the stability of the system.

A change in 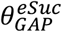 has direct effect on 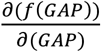, with higher 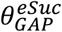 values increasing the effect that a change in the concentration of GAP has upon its own dynamics. Furthermore, 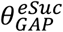 only affects 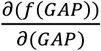, whereas a changes in other parameters of GAP also affect other entries in the Jacobian (e.g., 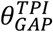 affects 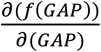 and 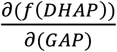. This could make GAP dynamics more resilient against effects of other metabolites and might constitute an overflow mechanism for excess carbon out of the CBC without influencing CBC dynamics. Similarly, an increase in 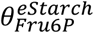 affects 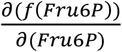, allowing more control over its own dynamics. However, as mentioned previously, the most crucial junction determining the stability was found to be at GAP and its various pathways. Thus, an increase in the absolute value of 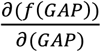 appears to be more beneficial than an absolute increase in 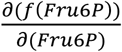.

### A shift in carbon partitioning towards sucrose synthesis increases metabolic stability

Based on the assumption that an increase in the absolute value of 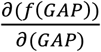 may stabilize the CBC, this suggests that an increase in the ratio of assimilated carbon going to sucrose biosynthesis rather than starch synthesis should increase the stability of the model as well. This would be a consequence of (Eq. 10)

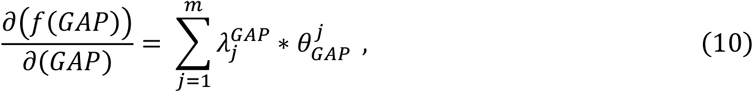

with 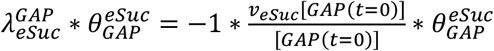. Thus, shifting the assimilated carbon towards sucrose synthesis rather than starch synthesis rate increased *v*_*eSuc*_ [*GAP*(*t* = 0)] and thus 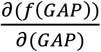 resulted in a gain in stability. To test this, the ratio of sucrose/starch allocation was shifted in the model and the results where again simulated 10^6^ times for each condition.

Supporting the hypothesis, results showed that an increased proportion of sucrose biosynthesis resulted in an increase in the stability of the system (Fig. 5). Interestingly, this effect was more pronounced under deficiency of photorespiration (24.4% *→* 46.2% stability vs. 79.8% *→* 86.1% stability, see Fig. 5).

**Fig. 5:**
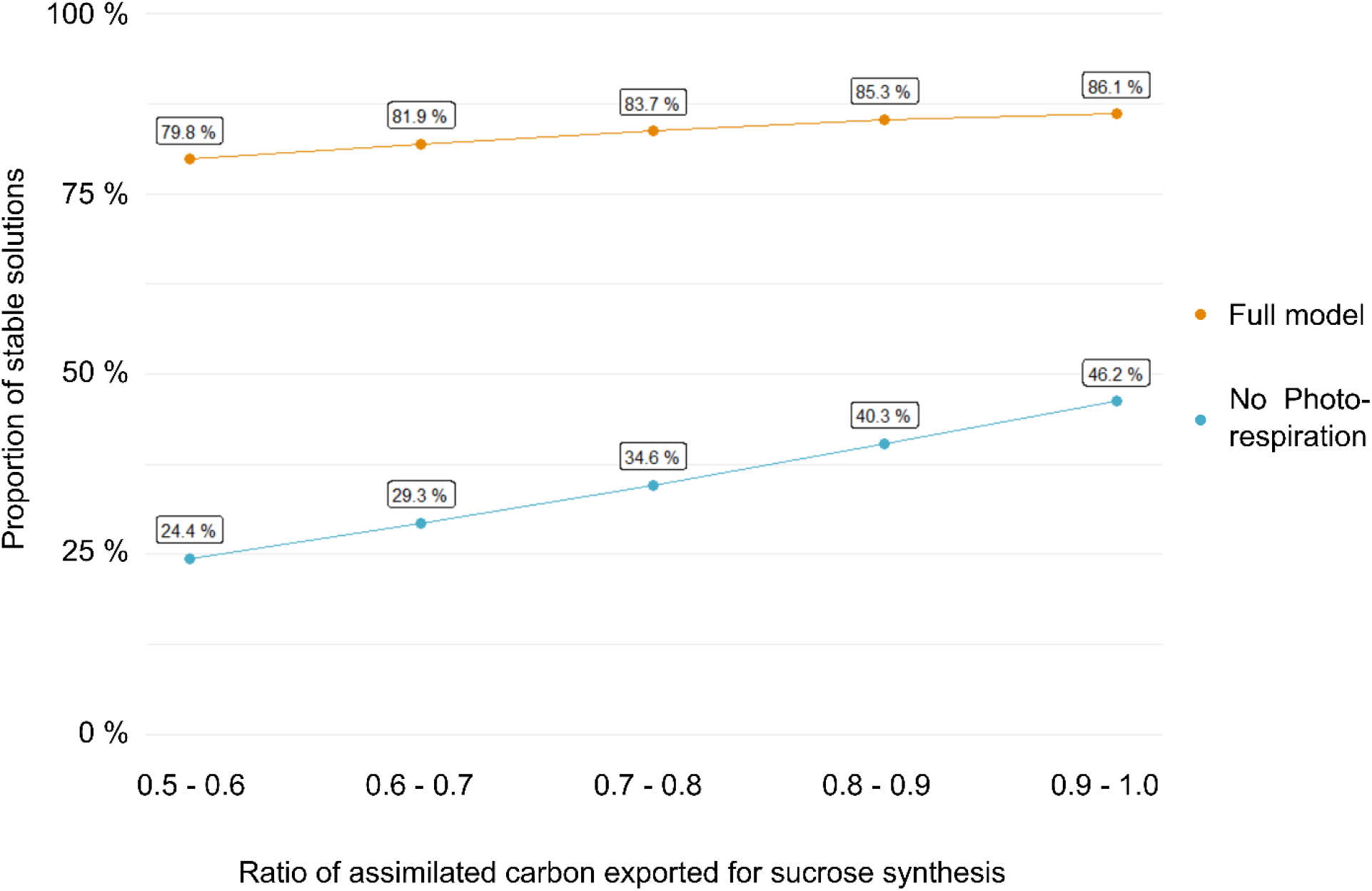
Portion of stable solutions with increasing carbon partitioning going towards sucrose synthesis. Each point was simulated 10^6^ times at different rations of sucrose synthesis randomized within an interval of 0.1 the ratio is given in C6-Sucrose/C6-Starch.

In addition to increased biosynthesis of soluble sugars, also accumulation of secondary metabolites is a well-known stress and acclimation response of plants, see e.g. (Winkel-Shirley, 2002; Doerfler et al., 2013). Following the finding that a shift form starch towards sucrose synthesis was able to stabilize the system, a model was created containing an Ery4P export to simulate the impact that induction of secondary metabolism via the shikimate pathway might have on the CBC. A graphical representation of this model, plus a model containing Ery4P export but no photorespiration, is provided in the supplements (Fig. S3). Similar as to what was observed for starch, the value of 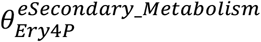 had no significant effect on the stability of the system (Table II).

**Table II:** Effect of 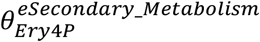 on the stability of the CBC.

| Model | Rank | Pearson Correlation | d<sub>n</sub> |
| --- | --- | --- | --- |
| Full model | 34 of 35 | -0.01 | 0.001 |
| No Photorespiration | 15 of 21 | 0.00 | 0.011 |

Based on this finding, we hypothesized that GAP export for sucrose synthesis is particularly able to stabilize the system which might result from its direct ability to control GAP dynamics. To test this hypothesis, the ratio of carbon going towards sucrose, starch or secondary metabolites was altered and proportion of stable solutions under each condition was quantified (Fig. 6).

**Fig. 6:**
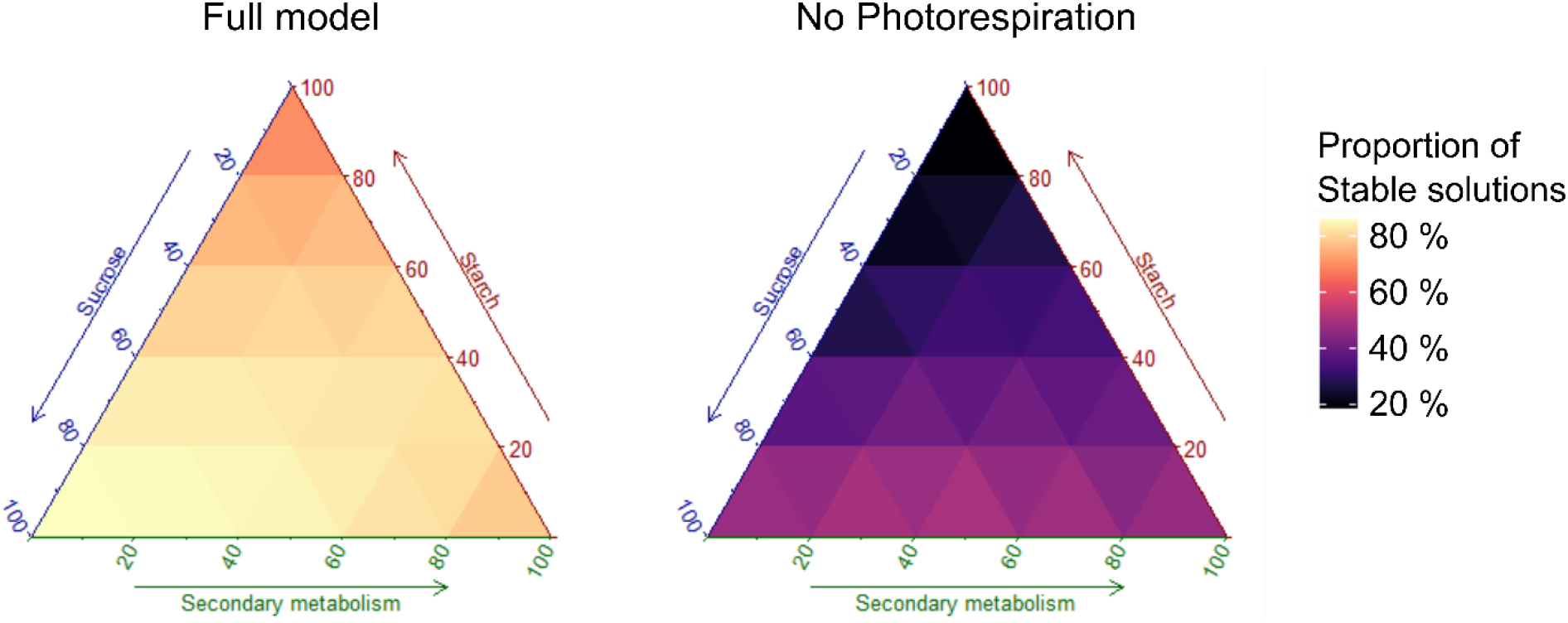
How carbon allocation affects the stability of the CBC. The ternary plot was constructed from 231 data points each simulated 10^4^ times per model. Arrows indicate the proportion of carbon flux in direction of sucrose, starch and/or secondary metabolites. Dark color indicates low proportion of stable solutions, light color indicates high proportion of stable solutions.

In presence of the photorespiratory pathway, carbon allocation towards sucrose proved to be the most effective in stabilizing the system (Fig. 6, left panel, full model). In contrast, a lack of photorespiration shifted most stable parameter combinations towards higher activity of secondary metabolism (40% - 60%). In both models, carbon channeling into starch biosynthesis contributed least to stability (0% - 20%). In summary, these results suggested that photorespiratory activity enables higher rates of sucrose biosynthesis without compromising the stability of the metabolic network. Further, the stabilizing effect of GAP export was only observed to be efficient if a sufficiently large parameter space for GAP export is present which might be achieved either through photorespiration or secondary metabolism (Fig. S4).

### The role of 2-PG in stabilizing the CBC

To analyze if regulation of the CBC by 2-PG though inhibition of TPI and/or SBPase may change the carbon partitioning towards a more stable system, a Branch and Bound (BnB) algorithm was applied with the purpose of searching for optimal strategy resulting in stable solutions. This analysis was challenged by the combinatorial problem of testing ∼4*10^12^ parameter combinations. Here, BnB enabled a computation time of, in total, < 700 hours (MATLAB® R2021a, Intel® Core™ i7-10700 @ 2.90 GHz).

In a situation where no enzymatic regulation, except for TPI and SBPase, was considered, both inhibition of TPI and SBPase by 2-PG lead to a decrease in stability (Fig. 7 A). This effect was strongest for TPI, where the proportion of stable models fell from 85.3% without inhibition to 6.1% with strong inhibition. Although less pronounced, a drop in stability from 85.3% to 80.6% could also be observed for SBPase. This drastically changed when considering full enzymatic regulation of the CBC, highlighting the importance of the BnB approach (Fig. 7 B). In 30% of near optimal solutions (no significant difference to the best solution, p > 0.01), a weak inhibition of TPI by 2-PG was observed. A weak inhibition by 2-PG of SBPase was observed in 15 % of all simulated models and strong inhibition in 75 % of solutions. This indicated a positive effect of regulation by 2-PG on the stability of the metabolic network with inhibition of SBPase taking a key role. Although this is more ambiguous for the inhibition of TPI, a considerable parameter space existed also for this inhibition (30% of all scenarios) which favors inhibition.

**Fig. 7:**
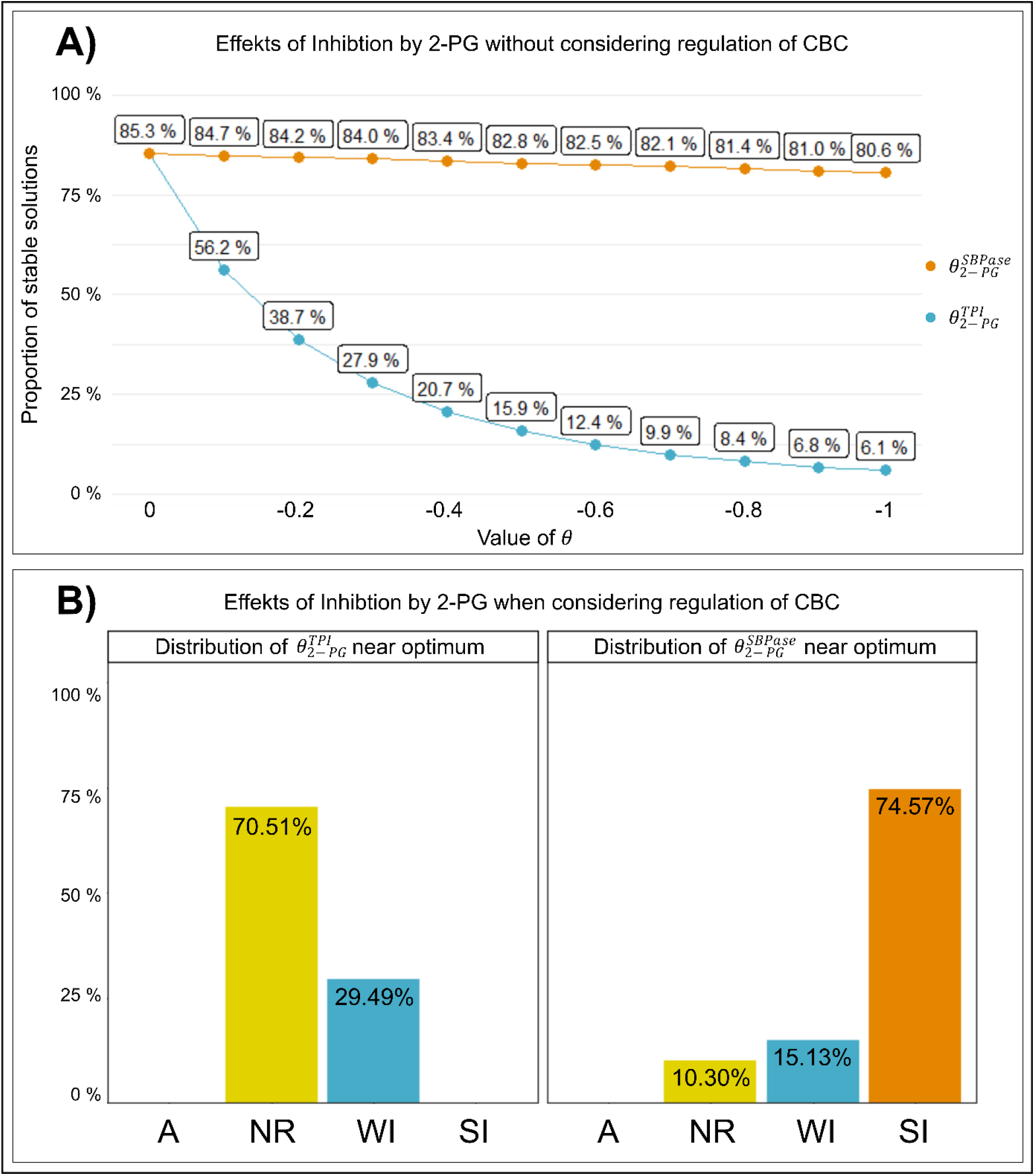
Effects of inhibition of TPI and SBPase by 2-PG. **A)** Change in stability with rising inhibition. Each point was simulated 10^5^ times. The blue line represents the effect of 2-PG on TPI. The orange line depicts the effect of 2-PG on SBPase. **B)** Results of parameter optimization considering regulation of the CBC. The bar plot depicts the distribution of regulatory interactions between 2-PG and TPI or SBPase found at the optimal solution and solutions which are not significantly worse (p > 0.01). A: activation (*θ* = 1), NR: no regulation (*θ* = 0), WI: weak inhibition (*θ* = −0.1), SI: strong inhibition (*θ* = −0.99).

## Discussion

Exposure to a sudden change of environmental conditions typically induces stress reactions in organisms to counteract and prevent irreversible damage of cells, tissues or organs. Stabilization of metabolism plays a central role in such stress response because it is preliminary to perceive and integrate environmental signals (Zhang et al., 2022). Stability properties of a biochemical reaction network can be quantified via an SKM approach (Steuer et al., 2006). With this approach, effects of single and/or multiple effectors and regulators on system stability can be estimated to yield non-intuitive information about metabolic pathways, pathway structures (Reznik and Segrè, 2010) or acclimation strategies (Fürtauer and Nägele, 2016). However, in metabolic systems, a vast variety of regulatory interactions between proteins and metabolites needs to be considered which rapidly results in extensive combinatorial problems for computation. To overcome some limitations of computation time, we applied a discreet parameter optimization approach. The importance of such an approach became evident when evaluating the effect of inhibition of CBC enzymes by 2-PG which finally suggested that photorespiration represents a trade-off between the carbon assimilation rate and the stability of cellular metabolism. This finding supplements and extends the previously suggested regulatory role of 2-PG for adjustment and allocation of chloroplast carbon flow (Flügel et al., 2017). The amount of 2-PG is tightly regulated by 2-PG phosphatase, PGLP, and its activity was shown to be essential for efficient carbon fixation and allocation (Flügel et al., 2017; Levey et al., 2019). Altogether, this suggests a strong impact of PGLP activity on stabilizing whole cell carbon metabolism after environmental perturbation. Further, simulations show that differential allocation of fixed carbon in either soluble carbohydrates, storage compounds or other (secondary) metabolism differentially affects stability of the whole CBC network. As a consequence, under sudden temperature and/or light changes, when ratios of Rubisco-driven carboxylation and oxygenation significantly deviate from the current homeostasis, metabolism needs to be reprogrammed to maximize stabilization capacities. The capability of re-stabilizing carbon fixation, and all downstream carbohydrate biosynthesis, is a critical variable in plant ecology and evolution because it significantly determines cell and tissue fate as well as growth and developmental success. Hence, combining biochemical and physiological experiments with stability analysis, e.g., by SKM, represents a promising approach to yield detailed insights into metabolic regulation. This information is essential to interpret experimental data on metabolite dynamics because underlying regulation might comprise non-intuitive patterns, e.g. metabolic cycles (Reznik and Segrè, 2010) or nested structures (Schaber et al., 2009) which limit interpretation of experimental findings. Further, it indicates the dependency of structural kinetic properties within a metabolic network which interconnects pathways across diverse subcellular compartments. We found that stabilization is achieved by increasing the parameter space of 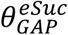 and 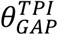 resulting in a stable solution (see Fig. 4). Owing to this and the noticeable correlation of 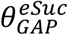 but not for 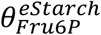, a shift from starch accumulation towards sucrose synthesis was able to stabilize the kinetic models (see Figs. 3 and 5). It has been described earlier that under stress, e.g., low temperature, carbon allocation is redirected from starch to sucrose biosynthesis (Strand et al., 1997; Strand et al., 1999). While this can be explained by the metabolic role of soluble carbohydrates as substrates for other pathways or osmotically active substances (Obata and Fernie, 2012), we hereby provide evidence for an additional role of this metabolic reprogramming in structural kinetic stabilization of carbon metabolism against environmentally induced perturbations.

Besides soluble carbohydrates, also secondary metabolites, e.g., flavonoids, are well-known to be involved in plant abiotic stress reactions, acclimation and tolerance mechanisms (Winkel-Shirley, 2002; Schulz et al., 2016; Naikoo et al., 2019). Yet, previously, evidence has also been provided for significant variation of extent of accumulation of secondary metabolites even across different natural accessions of *Arabidopsis thaliana* (Schulz et al., 2015). While significant correlation between tolerance measures, e.g., freezing tolerance, and flavonoid metabolism has been observed in these studies, some of the underlying mechanisms remain elusive. For example, transcript levels were found to be associated much stronger to freezing tolerance than absolute metabolite levels (Schulz et al., 2015). In context of findings of the present study this might highlight and support the hypothesis of metabolic stabilization by modification of efflux/influx capacities of the CBC and photorespiratory pathways towards starch, sucrose and secondary metabolism (see Fig. 6). While our analysis revealed, that in the full model partitioning of carbon towards sucrose biosynthesis improved the stability more than allocating carbon towards secondary metabolism, secondary metabolism was structurally able to provide a similar effect by increasing the admissive parameter space when oxygenation of RuBP was decreased. Finally, the optimal carbon partitioning was shifted under these conditions towards 40 – 50 % into secondary metabolism and 50 – 60 % into sucrose synthesis. Such a scenario might reflect a stress acclimated state of plant metabolism when rates of photorespiration are decreased again after initial stress response while sugars and secondary metabolites are significantly increased in their amount (Savitch et al., 2001; Doerfler et al., 2013). It further emphasizes the regulatory interaction between pathways with differential subcellular localization and indicates how stability may affect evolution of metabolism.

In conclusion, our study suggests a stabilizing role of photorespiration on the dynamics of cellular metabolism, thus representing a tradeoff between carbon assimilation and metabolic stability in a changing or fluctuating environment. Finally, shifting carbon partitioning from starch accumulation towards sucrose synthesis represents an effective measure for further increasing system stability, thus promoting our understanding of plant metabolic stress response.

## Supporting information

Matlab scripts

pseudocode BnB

## Acknowledgements

We would like to thank all members of Plant Evolutionary Cell Biology at the Faculty of Biology, LMU Munich, for many fruitful discussions. This work was funded by Deutsche Forschungsgemeinschaft, DFG (NA 1545/4-1).

## Supplementary legends

**Fig. S1: Graphical representation of models**. A) Full model. B) Model lacking photorespiration. C) Model with unspecific 2-PG efflux.

**Fig. S2: Parameters influencing the stability of the model lacking photorespiration**. A) Parameters ranked according to their influence on model stability. B) Most important reactions. Red = high importance, Yellow = moderate importance.

**Fig. S3: Graphical representation of models with export for secondary metabolism**. A) Full model with Ery4P export. B) Model lacking photorespiration with Ery4P export.

**Fig. S4: Effect of carbon partitioning between sucrose synthesis and secondary metabolism**. A) Top - Changes to model stability. B) Bottom - Effect on admissible parameter space.

