## Supplementary figures and images for "The trade-off function of photorespiration in a changing environment"

### Fig. S1

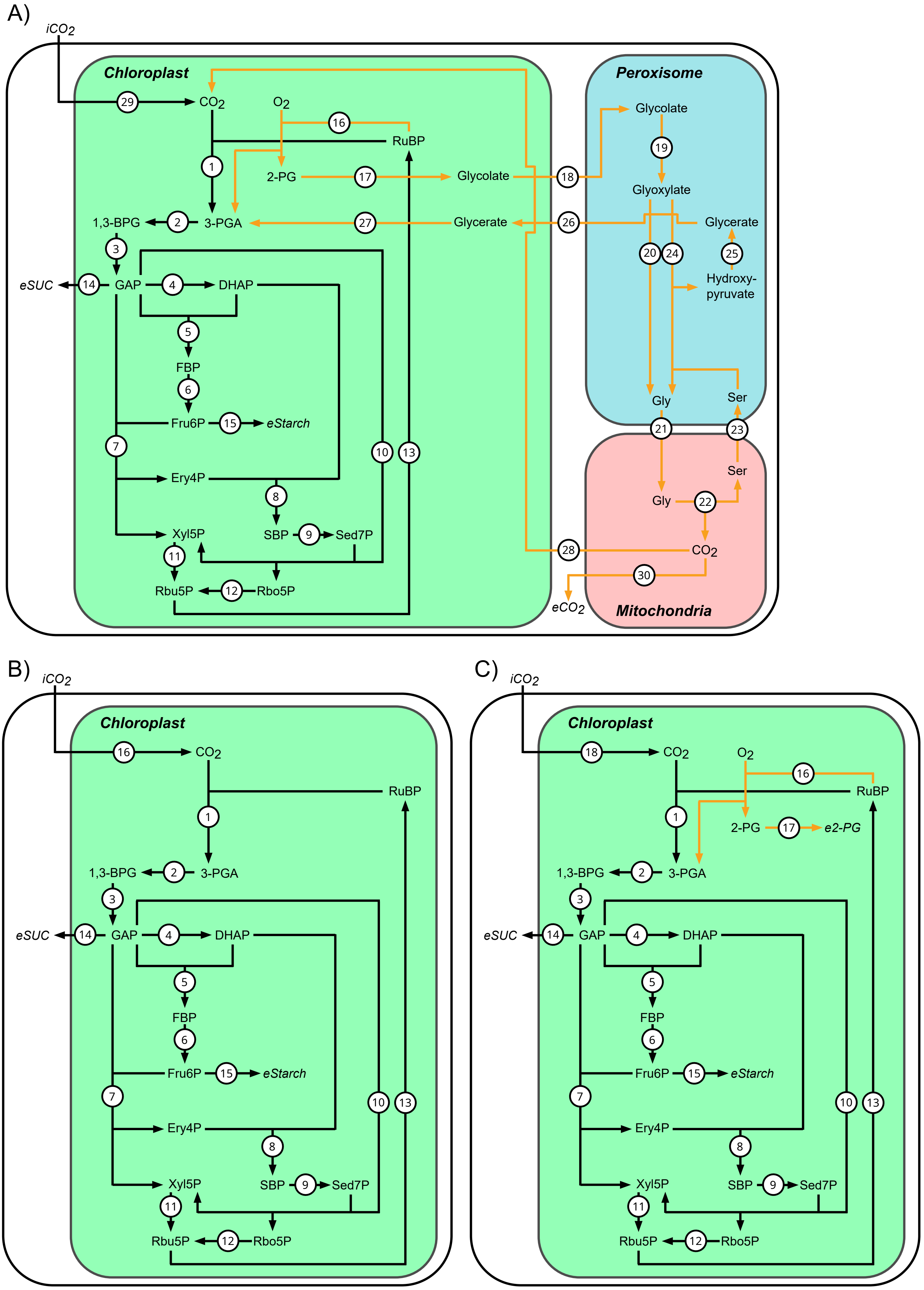

### Fig. S3

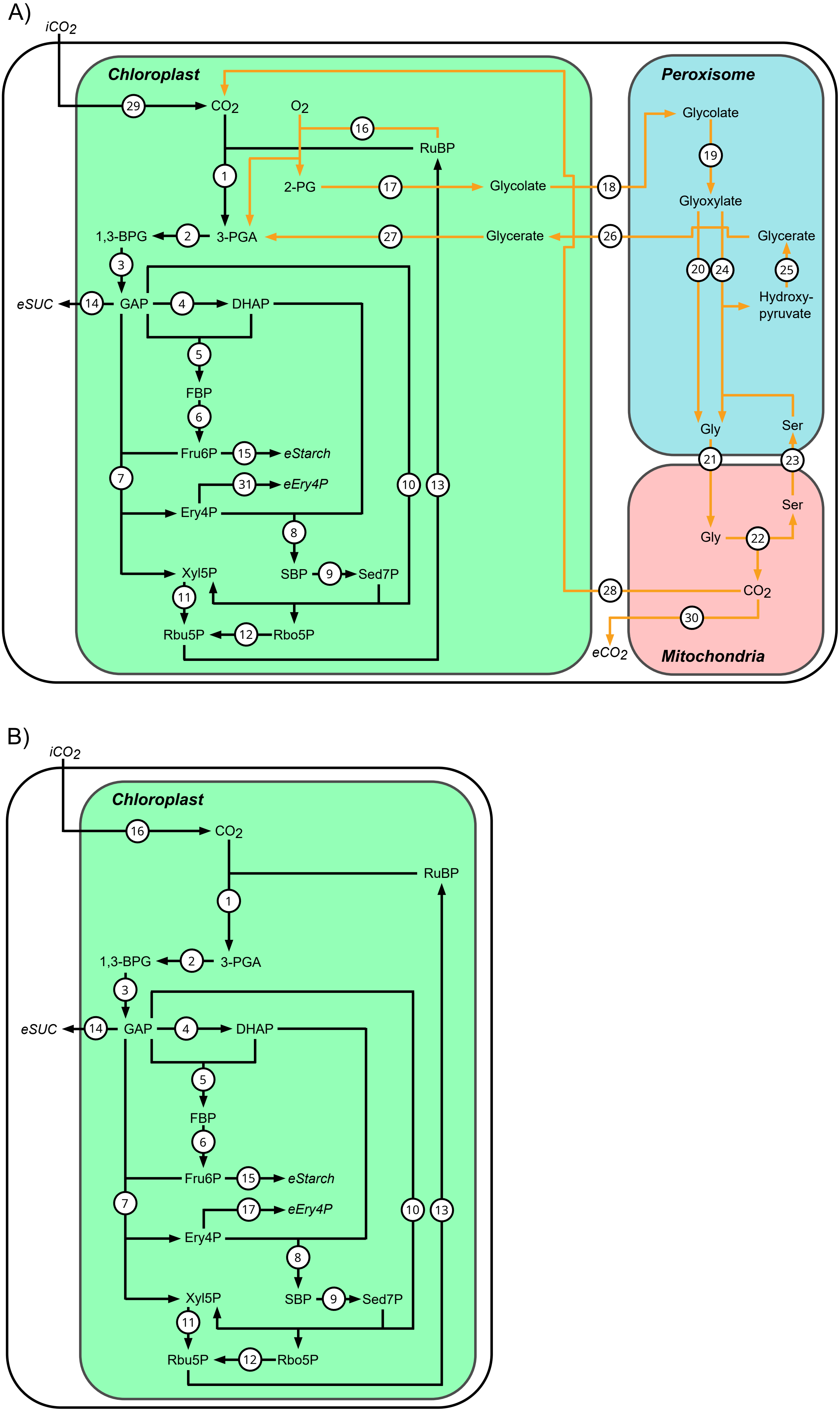

### Fig. S4

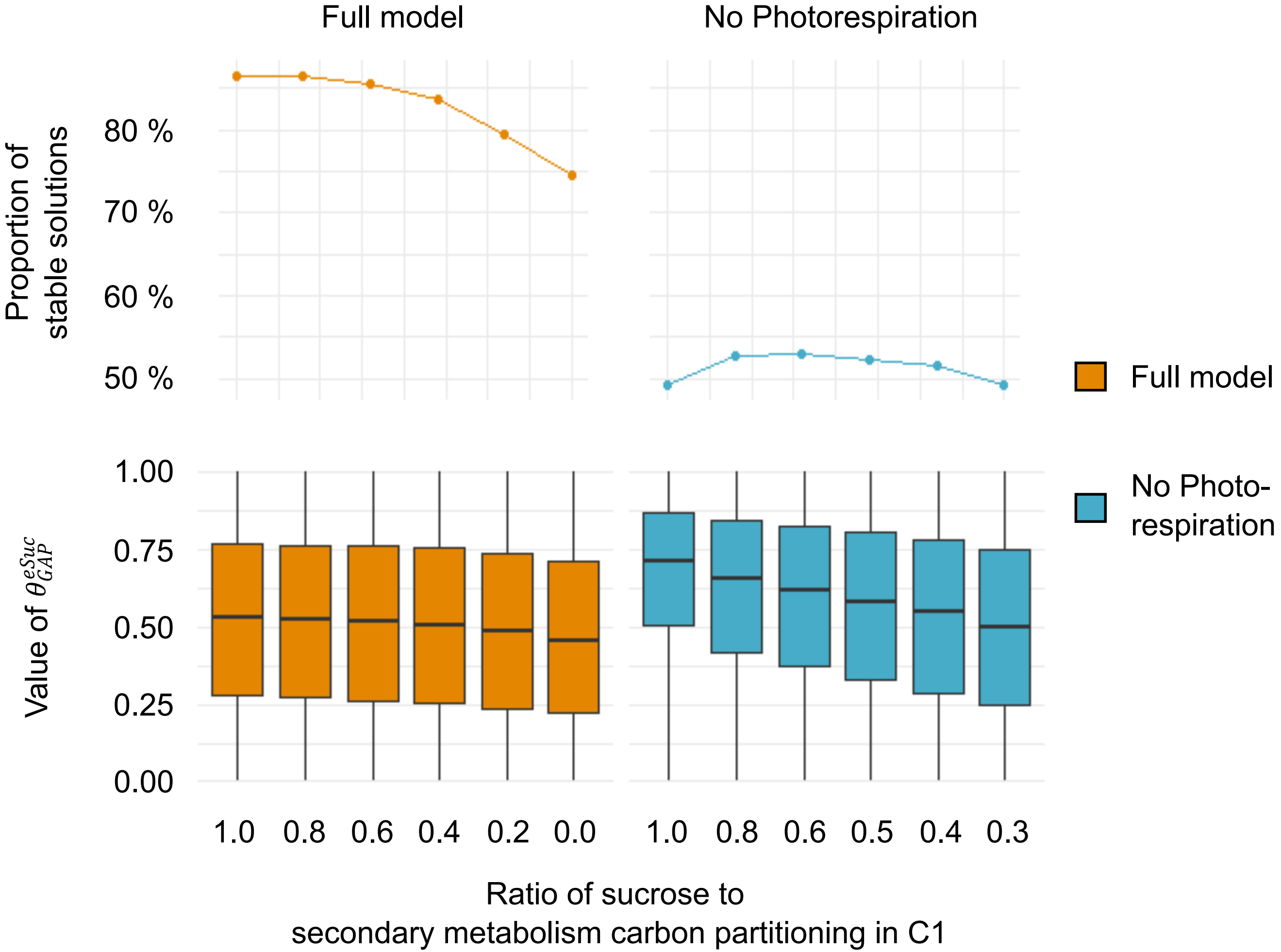
