## Supplementary material for "The trade-off function of photorespiration in a changing environment": Fig. S2

A)

| Overall rank | $\theta$ | *Pearson correlation* | *Rank of correlation* | $d_n = \|F_n - F_0\|$ | *Rank of $d_n$* | *Average of ranks* |
|---|---|---|---|---|---|---|
| *1* | $\theta_{GAP}^{TPI}$ | -0.07 | 2 | 0.2454 | 2 | 2.0 |
| *2* | $\theta_{GAP}^{eSuc}$ | -0.05 | 3 | 0.2892 | 1 | 2.0 |
| *3* | $\theta_{CO2}^{Rubisco}$ | -0.11 | 1 | 0.0102 | 5 | 3.0 |
| *4* | $\theta_{GAP}^{TK}$ | 0.04 | 5 | 0.1016 | 3 | 4.0 |
| *5* | $\theta_{RuBP}^{Rubisco}$ | 0.04 | 4 | 0.0028 | 7 | 5.5 |
| *6* | $\theta_{Fru6P}^{TK}$ | -0.01 | 10 | 0.0121 | 4 | 7.0 |
| *7* | $\theta_{3PGA}^{PGK}$ | 0.02 | 6 | 0.0021 | 11 | 8.5 |
| *8* | $\theta_{GAP}^{Ald}$ | -0.01 | 8 | 0.0026 | 9 | 8.5 |
| *9* | $\theta_{GAP}^{TK}$ | 0.01 | 7 | 0.0017 | 14 | 10.5 |
| *10* | $\theta_{Sed7P}^{TK}$ | 0.00 | 13 | 0.0027 | 8 | 10.5 |
| *11* | $\theta_{Ery4P}^{ALD}$ | 0.01 | 11 | 0.0017 | 13 | 12.0 |
| *12* | $\theta_{Fru6P}^{eStarch}$ | 0.00 | 20 | 0.0032 | 6 | 13.0 |

B)

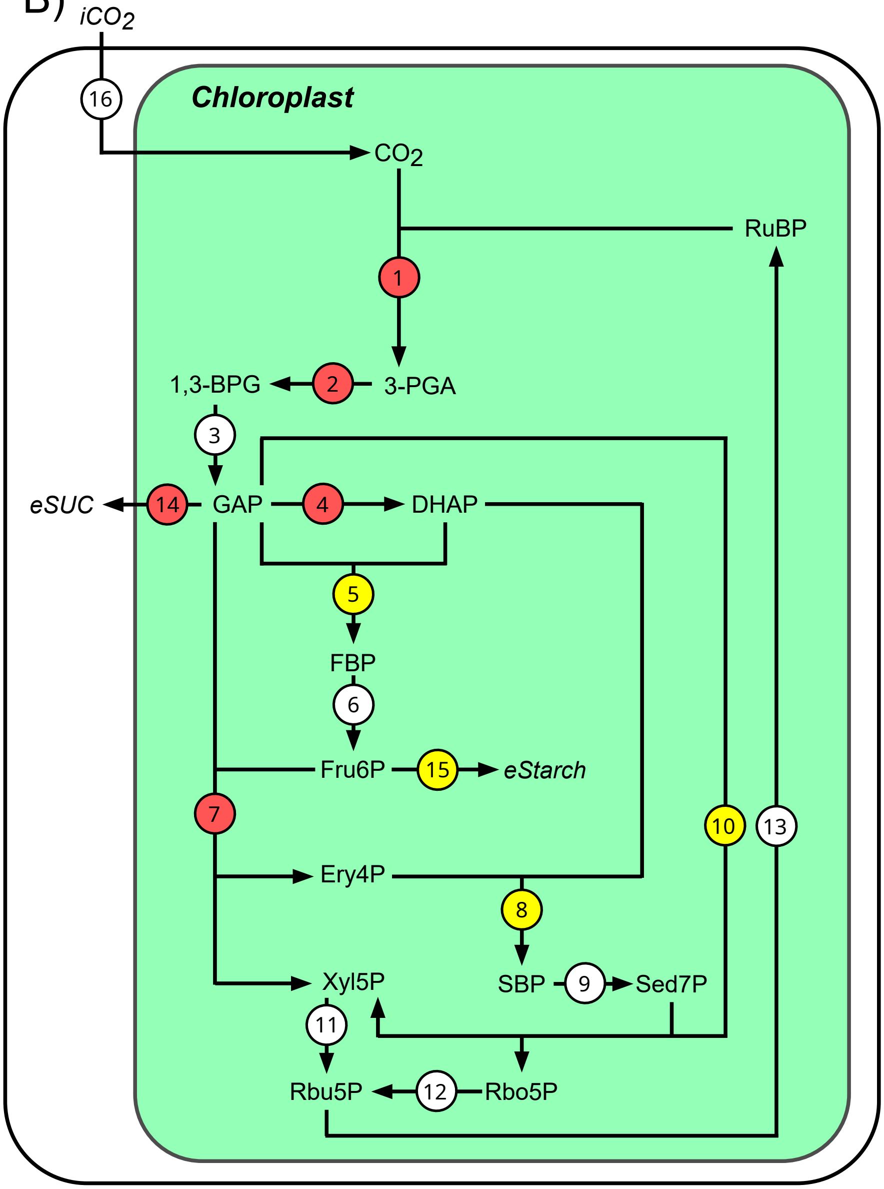
