## Supplementary material for "The trade-off function of photorespiration in a changing environment": Matlab scripts: Test_Stability_2_PG_efflux.m

```
%Specify number of iterations at start of run
numberruns = input('# of Replicates: ');

%In order to show Progress
x = numberruns/(numberruns/1000);

%Define storage for results

matrixdescription = {'F','D','Theta_1_13','Theta_1_14','Theta_2_1','Theta_3_2','Theta_4_3','Theta_5_3','Theta_5_4','Theta_6_5','Theta_7_3','Theta_7_6','Theta_8_4','Theta_8_7','Theta_9_8','Theta_10_3','Theta_10_9','Theta_11_11','Theta_12_10','Theta_13_12','Theta_14_3','Theta_15_6','Theta_16_13','Theta_17_14'};

Parameters = zeros(numberruns,length(matrixdescription));
result_eig = zeros(15,numberruns);
result_eigreal = zeros(15,numberruns);
    
%Get Model for Stichiometry

Phot_1 = IQMmodel('check_SS_2_PG_efflux.txt');
N = IQMstoichiometry(Phot_1);
    
for run_n = 1:numberruns,

    %Define Fluxes and Concentrationas
    
    F = 80 + 40 * rand(1); % rate of carboxylation of rubisco
    P = 0.2 + 0.2 * rand(1); % Prtion of Oxygenation to Carboxylaiton
    D = 0.8 + 0.1 * rand(1); % portion of assimilated carbon allocated to Sucrose synthesis. (1 - D) will go to Starch synthesis


    %specify reaction rates
    v = [F;
    2 *  F + P * F;
    2 * F + P * F;
    (2/3) * (F + P * F) + (1/2) * (1 - D) * (2 * F + P * F - (5/3) * (F + P * F));
    (1/3) * (F + P * F) + (1/2) * (1 - D) * (2 * F + P * F - (5/3) * (F + P * F));
    (1/3) * (F + P * F) + (1/2) * (1 - D) * (2 * F + P * F - (5/3) * (F + P * F));
    (1/3) * (F + P * F);
    (1/3) * (F + P * F);
    (1/3) * (F + P * F);
    (1/3) * (F + P * F);
    (2/3) * (F + P * F);
    (1/3) * (F + P * F);
    F + P * F;
    D * (2 * F + P * F - (5/3) * (F + P * F));
    (1/2) * (1 - D) * (2 * F + P * F - (5/3) * (F + P * F));
    P * F;
    P * F;
    F + P * F];
    
    %specify_starting_concentrations
    c = [0.59; %PGA
    0.001; %1,3-BPF
    0.01; %GAP
    0.27; %DHAP
    0.024; %FBP
    1.36; %Fru6P
    0.04; %Ery4P
    0.13; %SBP
    0.22; %Sed7P
    0.06; %Rbo5P
    0.04; %Xyl5P
    0.02; %Rbu5P
    0.14; %RuBP
    0.0018; %2-PG
    0.012]; %CO2

    %Generate Lambda

    LAMBDA = N .* v(:,ones(15,1))';
    LAMBDA=LAMBDA./c(:,ones(18,1));
    
    %Generate Theta
    
    Theta_1_13 = rand(1);
    Theta_1_14 = rand(1);
    Theta_2_1 = rand(1);
    Theta_3_2 = rand(1);
    Theta_4_3 = rand(1);
    Theta_5_3 = rand(1);
    Theta_5_4 = rand(1);
    Theta_6_5 = rand(1);
    Theta_7_3 = rand(1);
    Theta_7_6 = rand(1);
    Theta_8_4 = rand(1);
    Theta_8_7 = rand(1);
    Theta_9_8 = rand(1);
    Theta_10_3 = rand(1);
    Theta_10_9 = rand(1);
    Theta_11_11 = rand(1);
    Theta_12_10 = rand(1);
    Theta_13_12 = rand(1);
    Theta_14_3 = rand(1);
    Theta_15_6 = rand(1);
    Theta_16_13 = rand(1);
    Theta_17_14 = rand(1);
    

    THETA = [0,0,0,0,0,0,0,0,0,0,0,0,Theta_1_13,0,Theta_1_14;
    Theta_2_1,0,0,0,0,0,0,0,0,0,0,0,0,0,0;
    0,Theta_3_2,0,0,0,0,0,0,0,0,0,0,0,0,0;
    0,0,Theta_4_3,0,0,0,0,0,0,0,0,0,0,0,0;
    0,0,Theta_5_3,Theta_5_4,0,0,0,0,0,0,0,0,0,0,0;
    0,0,0,0,Theta_6_5,0,0,0,0,0,0,0,0,0,0;
    0,0,Theta_7_3,0,0,Theta_7_6,0,0,0,0,0,0,0,0,0;
    0,0,0,Theta_8_4,0,0,Theta_8_7,0,0,0,0,0,0,0,0;
    0,0,0,0,0,0,0,Theta_9_8,0,0,0,0,0,0,0;
    0,0,Theta_10_3,0,0,0,0,0,Theta_10_9,0,0,0,0,0,0;
    0,0,0,0,0,0,0,0,0,0,Theta_11_11,0,0,0,0;
    0,0,0,0,0,0,0,0,0,Theta_12_10,0,0,0,0,0;
    0,0,0,0,0,0,0,0,0,0,0,Theta_13_12,0,0,0;
    0,0,Theta_14_3,0,0,0,0,0,0,0,0,0,0,0,0;
    0,0,0,0,0,Theta_15_6,0,0,0,0,0,0,0,0,0;
    0,0,0,0,0,0,0,0,0,0,0,0,Theta_1_13,0,0;
    0,0,0,0,0,0,0,0,0,0,0,0,0,Theta_17_14,0;
    0,0,0,0,0,0,0,0,0,0,0,0,0,0,0];
    
    %Get Jacobian and eigenvalues

    J = LAMBDA * THETA;

    solution = eig(J);
    eigreal = real(solution);
    Parameters(run_n,:) = [F,D,Theta_1_13,Theta_1_14,Theta_2_1,Theta_3_2,Theta_4_3,Theta_5_3,Theta_5_4,Theta_6_5,Theta_7_3,Theta_7_6,Theta_8_4,Theta_8_7,Theta_9_8,Theta_10_3,Theta_10_9,Theta_11_11,Theta_12_10,Theta_13_12,Theta_14_3,Theta_15_6,Theta_16_13,Theta_17_14];
    result_eig(:,run_n) = solution;
    result_eigreal(:,run_n) = eigreal;
    result_max_eigreal = max(result_eigreal)';
    
    if rem(run_n,x) == 0
        disp(run_n)
    end
    
end

histogram(result_max_eigreal,13)
count = length(nonzeros(result_max_eigreal(result_max_eigreal<0)))

% Save Files (un-comment)

%writematrix(result_max_eigreal,'Max_eig.csv')
%writematrix(Parameters,'Parameters.csv')
%writematrix(result_eig','Eigenvalues.csv')
```
