## Supplementary material for "The trade-off function of photorespiration in a changing environment": Matlab scripts: BnB_SKM_CO2.m

```
% Branch and Bound Algorithem to find Heuristic and optimal solutions
% Using:
% DFS (Depth-first search) Search Strategy
% Wide Branching Strategy


clc;
clear;
close all;

disp('Starting BnB to find best regulation')

%% Load stoichiometric matrix

Phot_1 = IQMmodel('check_SS_v8_CO2.txt');
N = IQMstoichiometry(Phot_1);

%% Parameters of BnB

nVar = 21;                          % Specify number of Variables to Optimise
VarSize = [1 nVar];                 % Make Template Size

Discreat = [0, 1, -0.99, -0.1];    % Specify possible Values

disp(['Parameters set, maximum Number of final nodes: ' num2str(4^nVar)])

%% Initialization

PSOnum = 1;

% Loop to fix first variable in place and generate first noded for queue
for i = 1:4

    Disc.Position = 1;
    Disc.Value = Discreat(i);

    Node(i).Position = 1;
    Node(i).Value = Disc.Value;
    Node(i).Solution = PSO_BnB_CO2(Disc,N);
    
    disp(['Completed PSO number ' num2str(PSOnum) ' at Node Level ' num2str(Disc.Position) ' with ' num2str(100 * Node(i).Solution / 2000) '% Stable solutions'])
    PSOnum = PSOnum + 1;
        
end

% Establich framwork of solution structure and setting length to 0
SolvedNode(1).Solution = [];
SolvedNode(1).Mean = [];
SolvedNode(1).Value = [];
SolvedNode(1).Position = [];
SolvedNode(1) = [];

disp('Completed Initialization')

%% Mainloop

% main BnB loop
while isempty(Node) == 0

    Solution = [Node.Solution];
    [M,I] = max(Solution);

    x = zeros(VarSize);

    NodeLevel = Node(I).Position + 1;
    
    if NodeLevel < nVar
        
        PosNode = length(Node);
        
        for i = 1:4
            
            Disc.Position = [1 : NodeLevel];
            Disc.Value = [Node(I).Value Discreat(i)];

            Node(i + PosNode).Position = zeros(VarSize);
            Node(i + PosNode).Position = NodeLevel;
            Node(i + PosNode).Value = Disc.Value;
            Node(i + PosNode).Solution = PSO_BnB_CO2(Disc,N);
            
            disp(['Completed PSO number ' num2str(PSOnum) ' at Node Level ' num2str(NodeLevel) ' with ' num2str(100 * Node(i + PosNode).Solution / 2000) '% Stable solutions.'])
            PSOnum = PSOnum + 1;
            
        end
        
        Node(I) = [];
        disp(['The list has ' num2str(length(Node)) ' objects'])
        
        
    else
  
        PosSolvedNode = length(SolvedNode);

        for i = 1:4

            Disc.Position = [1 : NodeLevel];
            Disc.Value = [Node(I).Value Discreat(i)];
            x(Disc.Position) = Disc.Value;
            [Mean,Eig] = SKM_mean_CO2(x,false,N,0);
            SolvedNode(i + PosSolvedNode).Solution = Eig;
            SolvedNode(i + PosSolvedNode).Mean = Mean;
            SolvedNode(i + PosSolvedNode).Value = Disc.Value;
            SolvedNode(i + PosSolvedNode).Position = Disc.Position;
            
            disp(['Solved ' num2str(i + PosSolvedNode) ' Nodes at final Level'])

        end

        Node(I) = [];
        SolvedSolution = [SolvedNode.Solution];
        BestSol = max(SolvedSolution);
        Solution = [Node.Solution];
        if BestSol/2000 < 0.995
           
            Prune = find(2.33 < ((BestSol/2000) - (Solution/2000)) ./ sqrt(((BestSol + Solution)/4000) .* (1-((BestSol + Solution)/4000))*(1/1000)));
            
        else
            
            Prune = find(Solution/2000 < 0.995);
            
        end
        Node(Prune) = [];

    end

end


%% Results

Result = rmfield(SolvedNode,'Position');
Result_A = struct2array(Result);
Result_M = reshape(Result_A,nVar + 2,[])';
writematrix(Result_M,'Result_BnB.csv')

disp(['Competed parameter search: ' num2str(100 * (1 - (length(SolvedNode)/4^nVar))) '% of final Nodes where pruned.'])
```
