## Supplementary material for "The trade-off function of photorespiration in a changing environment": Matlab scripts: PSO_BnB_CO2.m

```
function BestCostOpti = PSO_BnB_CO2(Disc,N)

    %% Problem Definition

    CostFunction = @(x,GB_P) SKM_CO2(x,false,N,GB_P); %Costfunction

    nVar = 21;               % Specify number of Variables to Optimise

    VarSize = [1 nVar];      % Make Template Size

    VarMin = -0.99;          % Lower Bond of Variables - strong inhibition
    VarMax =  1;             % Upper Bond of Variables - strong activation

    %% Prameters of PSO

    kappa = 1;
    phi1 = 2.05;
    phi2 = 2.05;
    phi = phi1 + phi2;
    chi = 2*kappa/abs(2-phi-sqrt(phi^2-4*phi));


    MaxIt = 1000;                 % Maximum Number of Iterations
    nPop = 35;                    % Population Size (Swarm Size)
    w = chi;                      % Inertia Coefficient
    c1 = phi1*chi;                % Personal Acceleration Coefficient
    c2 = phi2*chi;                % Social Acceleration Coefficient

    MaxValocity = 0.2 * (VarMax - VarMin);
    MinValocity = -MaxValocity;

    %% Initialization

    empty_particle.Position = [];
    empty_particle.Velocity = [];
    empty_particle.Cost = [];
    empty_particle.Best.Position = [];
    empty_particle.Best.Cost = [];

    particle = repmat(empty_particle, nPop, 1);

    GlobalBest.Cost = -inf;

    for i = 1:nPop

        particle(i).Position = unifrnd(VarMin, VarMax, VarSize);
        
        particle(i).Position(1,50:nVar) = 0;
        particle(i).Position = particle(i).Position(randperm(length(particle(i).Position)));
        particle(i).Position(Disc.Position) = Disc.Value;

        particle(i).Velocity = zeros(VarSize);

        particle(i).Cost = CostFunction(particle(i).Position,0);

        particle(i).Best.Position = particle(i).Position;
        particle(i).Best.Cost = particle(i).Cost;

        if particle(i).Best.Cost > GlobalBest.Cost
            GlobalBest = particle(i).Best;
        end

    end

    BestCosts = zeros(MaxIt, 1);

    %% Main Loop of PSO

    for it = 1:MaxIt

        for i=1:nPop

            particle(i).Velocity = w*particle(i).Velocity ...
                + c1*rand(VarSize).*(particle(i).Best.Position - particle(i).Position) ...
                + c2*rand(VarSize).*(GlobalBest.Position - particle(i).Position);

            particle(i).Velocity = max(particle(i).Velocity, MinValocity);
            particle(i).Velocity = min(particle(i).Velocity, MaxValocity);

            particle(i).Position = particle(i).Position + particle(i).Velocity;
            particle(i).Position(Disc.Position) = Disc.Value;

            particle(i).Position = max(particle(i).Position, VarMin);
            particle(i).Position = min(particle(i).Position, VarMax);

            particle(i).Cost = CostFunction(particle(i).Position,GlobalBest.Cost);

            if particle(i).Cost > particle(i).Best.Cost

                particle(i).Best.Cost = particle(i).Cost;
                particle(i).Best.Position = particle(i).Position;

                if particle(i).Best.Cost > GlobalBest.Cost
                     GlobalBest = particle(i).Best;
                end

            end

        end

        % Store Best Cost
        BestCosts(it) = GlobalBest.Cost;
        
        disp(['Iteration ' num2str(it) ': Stable solutions = ' num2str(100*BestCosts(it)/2000) '%']);

        if BestCosts(it)/2000 > 0.995
            break
        end

        if it > 30 && ((BestCosts(it)/2000) - (BestCosts(it-30)/2000)) / sqrt(((BestCosts(it) ...
                + BestCosts(it-30))/4000) * (1-((BestCosts(it) + BestCosts(it-30))/4000))*(1/1000)) < 1.64
            break
        end

    end
    BestCostOpti = BestCosts(it);
end
    
```
