## Supplementary material for "The trade-off function of photorespiration in a changing environment": Matlab scripts: SKM_CO2.m

```
function Eig = SKM_CO2(x,ShowPlot,N,GB)

    %Specify number of iterations at start of run
    numberruns = 2000;

    %Define storage for results
    
    result_eigreal = zeros(26,numberruns);

    for run_n = 1:numberruns

        %Define Fluxes and Concentrationas. Phi = P (0.2|0.4), Lambda = L (0|0.5), Alpha
        %= A (0.75|0.95)

        F = 80 + 40 * rand(1); % rate of carboxylation of rubisco
        P = 0.2 + 0.2 * rand(1); % Prtion of Oxygenation to Carboxylaiton
        D = 0.8 + 0.1 * rand(1); % portion of assimilated carbon allocated to Sucrose synthesis. (1 - D) will go to Starch synthesis
        A = 0 + 1 * rand(1); % portion of photorespiratory reassimilated


        %specify reaction rates
        v = [F;
        2 * F + P * F + (1/2) * P * F;
        2 * F + P * F + (1/2) * P * F;
        (2/3) * (F + P * F) + (1/2) * (1 - D) * (2 * F + P * F + (1/2) * P * F - (5/3) * (F + P * F));
        (1/3) * (F + P * F) + (1/2) * (1 - D) * (2 * F + P * F + (1/2) * P * F - (5/3) * (F + P * F));
        (1/3) * (F + P * F) + (1/2) * (1 - D) * (2 * F + P * F + (1/2) * P * F - (5/3) * (F + P * F));
        (1/3) * (F + P * F);
        (1/3) * (F + P * F);
        (1/3) * (F + P * F);
        (1/3) * (F + P * F);
        (2/3) * (F + P * F);
        (1/3) * (F + P * F);
        F + P * F;
        D * (2 * F + P * F + (1/2) * P * F - (5/3) * (F + P * F));
        (1/2) * (1 - D) * (2 * F + P * F + (1/2) * P * F - (5/3) * (F + P * F));
        P * F;
        P * F;
        P * F;
        P * F;
        (1/2) * P * F;
        P * F;
        (1/2) * P * F;
        (1/2) * P * F;
        (1/2) * P * F;
        (1/2) * P * F;
        (1/2) * P * F;
        (1/2) * P * F;
        A * (1/2) * P * F;
        F - A * (1/2) * P * F;
        (1 - A) * (1/2) * P * F];

        %specify_starting_concentrations
        c = [0.59; %PGA
        0.001; %1,3-BPF
        0.01; %GAP
        0.27; %DHAP
        0.024; %FBP
        1.36; %Fru6P
        0.04; %Ery4P
        0.13; %SBP
        0.22; %Sed7P
        0.06; %Rbo5P
        0.04; %Xyl5P
        0.02; %Rbu5P
        0.14; %RuBP
        0.0018; %2-PG
        0.2; %Glycolate_chl
        0.1; %Glycolate_per
        0.3; %Glyoxylate
        0.49; %Gly_per
        0.57; %Gly_mit
        23.1; %Ser_mit
        3.4; %Ser_per
        0.23; %Hydroxypyruvate
        0.14; %Glycerate_per
        0.289; %Glycerate_chl
        0.012;
        0.012];        

        %Generate Lambda

        LAMBDA = N .* v(:,ones(26,1))';
        LAMBDA=LAMBDA./c(:,ones(30,1));

        %Generate Theta

        Theta = rand(1,34);
    
        THETA = [x(1),0,0,0,x(2),0,0,x(3),0,0,0,0,Theta(1),0,0,0,0,0,0,0,0,0,0,0,Theta(2),0;
        Theta(3),x(4),0,0,0,0,0,0,0,0,0,0,0,0,0,0,0,0,0,0,0,0,0,0,0,0;
        0,Theta(4),0,0,0,0,0,0,0,0,0,0,0,0,0,0,0,0,0,0,0,0,0,0,0,0;
        0,0,Theta(5),x(5),0,0,0,0,0,0,0,0,0,x(6),0,0,0,0,0,0,0,0,0,0,0,0;
        0,0,Theta(6),Theta(7),0,0,0,0,0,0,0,0,0,0,0,0,0,0,0,0,0,0,0,0,0,0;
        0,0,0,0,Theta(8),x(12),0,0,0,0,0,0,0,0,0,0,0,0,0,0,0,0,0,0,0,0;
        0,0,Theta(9),0,0,Theta(10),x(14),0,x(15),x(16),x(17),0,0,0,0,0,0,0,0,0,0,0,0,0,0,0;
        0,0,0,Theta(11),0,0,Theta(12),0,0,0,0,0,0,0,0,0,0,0,0,0,0,0,0,0,0,0;
        0,0,0,0,0,0,0,Theta(13),x(7),0,0,0,0,x(8),0,0,0,0,0,0,0,0,0,0,0,0;
        0,0,Theta(14),0,0,x(18),x(19),0,Theta(15),x(20),x(21),0,0,0,0,0,0,0,0,0,0,0,0,0,0,0;
        0,0,0,0,0,0,0,0,0,0,Theta(16),0,0,0,0,0,0,0,0,0,0,0,0,0,0,0;
        0,0,0,0,0,0,0,0,0,Theta(17),0,0,0,0,0,0,0,0,0,0,0,0,0,0,0,0;
        x(13),0,0,0,0,0,0,0,0,0,0,Theta(18),0,0,0,0,0,0,0,0,0,0,0,0,0,0;
        0,0,Theta(19),0,0,0,0,0,0,0,0,0,0,0,0,0,0,0,0,0,0,0,0,0,0,0;
        0,0,0,0,0,Theta(20),0,0,0,0,0,0,0,0,0,0,0,0,0,0,0,0,0,0,0,0;
        x(1),0,0,0,x(2),0,0,x(3),0,0,0,0,Theta(1),0,0,0,0,0,0,0,0,0,0,0,0,0;
        0,0,0,0,0,0,0,0,0,0,0,0,0,Theta(21),0,0,0,0,0,0,0,0,0,0,0,0;
        0,0,0,0,0,0,0,0,0,0,0,0,0,0,Theta(22),0,0,0,0,0,0,0,0,0,0,0;
        0,0,0,0,0,0,0,0,0,0,0,0,0,0,0,Theta(23),0,0,0,0,0,0,0,0,0,0;
        0,0,0,0,0,0,0,0,0,0,0,0,0,0,0,0,Theta(24),x(9),0,0,0,0,0,0,0,0;
        0,0,0,0,0,0,0,0,0,0,0,0,0,0,0,0,0,Theta(25),0,0,0,0,0,0,0,0;
        0,0,0,0,0,0,0,0,0,0,0,0,0,0,0,0,0,0,Theta(26),0,0,0,0,0,0,0;
        0,0,0,0,0,0,0,0,0,0,0,0,0,0,0,0,0,0,0,Theta(27),0,0,0,0,0,0;
        0,0,0,0,0,0,0,0,0,0,0,0,0,0,0,0,Theta(28),x(10),0,0,Theta(29),x(11),0,0,0,0;
        0,0,0,0,0,0,0,0,0,0,0,0,0,0,0,0,0,0,0,0,0,Theta(30),0,0,0,0;
        0,0,0,0,0,0,0,0,0,0,0,0,0,0,0,0,0,0,0,0,0,0,Theta(31),0,0,0;
        0,0,0,0,0,0,0,0,0,0,0,0,0,0,0,0,0,0,0,0,0,0,0,Theta(32),0,0;
        0,0,0,0,0,0,0,0,0,0,0,0,0,0,0,0,0,0,0,0,0,0,0,0,0,Theta(33);
        0,0,0,0,0,0,0,0,0,0,0,0,0,0,0,0,0,0,0,0,0,0,0,0,0,0;
        0,0,0,0,0,0,0,0,0,0,0,0,0,0,0,0,0,0,0,0,0,0,0,0,0,Theta(34)];

        %Get Jacobian and eigenvalues

        J = LAMBDA * THETA;

        solution = eig(J);
        eigreal = real(solution);
        result_eigreal(:,run_n) = eigreal;
        
        result_max_eigreal = max(result_eigreal)';
        Res = length(nonzeros(result_max_eigreal(result_max_eigreal<0)));
        p = Res/run_n;
        upper_con_int = p + 2.58*sqrt(p*(1-p)/run_n);
        
        if run_n > 100 && upper_con_int < GB/2000
            break
        end

    end
    
    result_max_eigreal = max(result_eigreal)';
    Eig = length(nonzeros(result_max_eigreal(result_max_eigreal<0)));
    
    if ShowPlot == true
        histogram(result_max_eigreal,13)
    end
    
end
```
