## Supplementary material for "The trade-off function of photorespiration in a changing environment": Matlab scripts: Test_Stability_full_model.m

```
clc;
clear;
close all;

%Specify number of iterations at start of run
numberruns = input('# of Replicates: ');

%In order to show Progress
x = numberruns/(numberruns/1000);

%Define storage for results

matrixdescription = {'F','P','D','A','Theta_1_13','Theta_2_1','Theta_3_2','Theta_4_3','Theta_5_3','Theta_5_4','Theta_6_5','Theta_7_3','Theta_7_6','Theta_8_4','Theta_8_7','Theta_9_8','Theta_10_3','Theta_10_9','Theta_11_11','Theta_12_10','Theta_13_12','Theta_14_3','Theta_15_6','Theta_17_14','Theta_18_15','Theta_19_16','Theta_20_17','Theta_21_18','Theta_22_19','Theta_23_20','Theta_24_17','Theta_24_21','Theta_25_22','Theta_26_23','Theta_27_24','Theta_1_25','Theta_28_26','Theta_30_26'};

Parameters = zeros(numberruns,length(matrixdescription));
result_eig = zeros(26,numberruns);
result_eigreal = zeros(26,numberruns);
    
%Get Model for Stichiometry

Phot_1 = IQMmodel('check_SS_full_model.txt');
N = IQMstoichiometry(Phot_1);
    
parfor run_n = 1:numberruns

    %Define Fluxes and Concentrationas
    
    F = 80 + 40 * rand(1); % rate of carboxylation of rubisco
    P = 0.2 + 0.2 * rand(1); % Prtion of Oxygenation to Carboxylaiton
    D = 0.8 + 0.1 * rand(1); % portion of assimilated carbon allocated to Sucrose synthesis. (1 - D) will go to Starch synthesis
    A = 0 + 1 * rand(1); % portion of photorespiratory reassimilated


    %specify reaction rates
    v = [F;
    2 * F + P * F + (1/2) * P * F;
    2 * F + P * F + (1/2) * P * F;
    (2/3) * (F + P * F) + (1/2) * (1 - D) * (2 * F + P * F + (1/2) * P * F - (5/3) * (F + P * F));
    (1/3) * (F + P * F) + (1/2) * (1 - D) * (2 * F + P * F + (1/2) * P * F - (5/3) * (F + P * F));
    (1/3) * (F + P * F) + (1/2) * (1 - D) * (2 * F + P * F + (1/2) * P * F - (5/3) * (F + P * F));
    (1/3) * (F + P * F);
    (1/3) * (F + P * F);
    (1/3) * (F + P * F);
    (1/3) * (F + P * F);
    (2/3) * (F + P * F);
    (1/3) * (F + P * F);
    F + P * F;
    D * (2 * F + P * F + (1/2) * P * F - (5/3) * (F + P * F));
    (1/2) * (1 - D) * (2 * F + P * F + (1/2) * P * F - (5/3) * (F + P * F));
    P * F;
    P * F;
    P * F;
    P * F;
    (1/2) * P * F;
    P * F;
    (1/2) * P * F;
    (1/2) * P * F;
    (1/2) * P * F;
    (1/2) * P * F;
    (1/2) * P * F;
    (1/2) * P * F;
    A * (1/2) * P * F;
    F - A * (1/2) * P * F;
    (1 - A) * (1/2) * P * F];
    
    %specify_starting_concentrations
    c = [1.59; %PGA
    0.001; %1,3-BPF
    0.01; %GAP
    0.27; %DHAP
    0.024; %FBP
    1.36; %Fru6P
    0.04; %Ery4P
    0.13; %SBP
    0.22; %Sed7P
    0.06; %Rbo5P
    0.04; %Xyl5P
    0.02; %Rbu5P
    0.14; %RuBP
    0.0018; %2-PG
    0.2; %Glycolate_chl
    0.1; %Glycolate_per
    0.3; %Glyoxylate
    0.49; %Gly_per
    0.57; %Gly_mit
    23.1; %Ser_mit
    3.4; %Ser_per
    0.23; %Hydroxypyruvate
    0.14; %Glycerate_per
    0.289; %Glycerate_chl
    0.012; %CO2 
    0.012]; %CO2_mit

    %Generate Lambda

    LAMBDA = N .* v(:,ones(26,1))';
    LAMBDA=LAMBDA./c(:,ones(30,1));
    
    %Generate Theta
    
    Theta_1_13 = rand(1);
    Theta_2_1 = rand(1);
    Theta_3_2 = rand(1);
    Theta_4_3 = rand(1);
    Theta_5_3 = rand(1);
    Theta_5_4 = rand(1);
    Theta_6_5 = rand(1);
    Theta_7_3 = rand(1);
    Theta_7_6 = rand(1);
    Theta_8_4 = rand(1);
    Theta_8_7 = rand(1);
    Theta_9_8 = rand(1);
    Theta_10_3 = rand(1);
    Theta_10_9 = rand(1);
    Theta_11_11 = rand(1);
    Theta_12_10 = rand(1);
    Theta_13_12 = rand(1);
    Theta_14_3 = rand(1);
    Theta_15_6 = rand(1);
    Theta_17_14 = rand(1);
    Theta_18_15 = rand(1);
    Theta_19_16 = rand(1);
    Theta_20_17 = rand(1);
    Theta_21_18 = rand(1);
    Theta_22_19 = rand(1);
    Theta_23_20 = rand(1);
    Theta_24_17 = rand(1);
    Theta_24_21 = rand(1);
    Theta_25_22 = rand(1);
    Theta_26_23 = rand(1);
    Theta_27_24 = rand(1);
    Theta_1_25 = rand(1);
    Theta_28_26 = rand(1);
    Theta_30_26 = rand(1);
    

    THETA = [0,0,0,0,0,0,0,0,0,0,0,0,Theta_1_13,0,0,0,0,0,0,0,0,0,0,0,Theta_1_25,0;
    Theta_2_1,0,0,0,0,0,0,0,0,0,0,0,0,0,0,0,0,0,0,0,0,0,0,0,0,0;
    0,Theta_3_2,0,0,0,0,0,0,0,0,0,0,0,0,0,0,0,0,0,0,0,0,0,0,0,0;
    0,0,Theta_4_3,0,0,0,0,0,0,0,0,0,0,0,0,0,0,0,0,0,0,0,0,0,0,0;
    0,0,Theta_5_3,Theta_5_4,0,0,0,0,0,0,0,0,0,0,0,0,0,0,0,0,0,0,0,0,0,0;
    0,0,0,0,Theta_6_5,0,0,0,0,0,0,0,0,0,0,0,0,0,0,0,0,0,0,0,0,0;
    0,0,Theta_7_3,0,0,Theta_7_6,0,0,0,0,0,0,0,0,0,0,0,0,0,0,0,0,0,0,0,0;
    0,0,0,Theta_8_4,0,0,Theta_8_7,0,0,0,0,0,0,0,0,0,0,0,0,0,0,0,0,0,0,0;
    0,0,0,0,0,0,0,Theta_9_8,0,0,0,0,0,-0.9,0,0,0,0,0,0,0,0,0,0,0,0;
    0,0,Theta_10_3,0,0,0,0,0,Theta_10_9,0,0,0,0,0,0,0,0,0,0,0,0,0,0,0,0,0;
    0,0,0,0,0,0,0,0,0,0,Theta_11_11,0,0,0,0,0,0,0,0,0,0,0,0,0,0,0;
    0,0,0,0,0,0,0,0,0,Theta_12_10,0,0,0,0,0,0,0,0,0,0,0,0,0,0,0,0;
    0,0,0,0,0,0,0,0,0,0,0,Theta_13_12,0,0,0,0,0,0,0,0,0,0,0,0,0,0;
    0,0,Theta_14_3,0,0,0,0,0,0,0,0,0,0,0,0,0,0,0,0,0,0,0,0,0,0,0;
    0,0,0,0,0,Theta_15_6,0,0,0,0,0,0,0,0,0,0,0,0,0,0,0,0,0,0,0,0;
    0,0,0,0,0,0,0,0,0,0,0,0,Theta_1_13,0,0,0,0,0,0,0,0,0,0,0,0,0;
    0,0,0,0,0,0,0,0,0,0,0,0,0,Theta_17_14,0,0,0,0,0,0,0,0,0,0,0,0;
    0,0,0,0,0,0,0,0,0,0,0,0,0,0,Theta_18_15,0,0,0,0,0,0,0,0,0,0,0;
    0,0,0,0,0,0,0,0,0,0,0,0,0,0,0,Theta_19_16,0,0,0,0,0,0,0,0,0,0;
    0,0,0,0,0,0,0,0,0,0,0,0,0,0,0,0,Theta_20_17,0,0,0,0,0,0,0,0,0;
    0,0,0,0,0,0,0,0,0,0,0,0,0,0,0,0,0,Theta_21_18,0,0,0,0,0,0,0,0;
    0,0,0,0,0,0,0,0,0,0,0,0,0,0,0,0,0,0,Theta_22_19,0,0,0,0,0,0,0;
    0,0,0,0,0,0,0,0,0,0,0,0,0,0,0,0,0,0,0,Theta_23_20,0,0,0,0,0,0;
    0,0,0,0,0,0,0,0,0,0,0,0,0,0,0,0,Theta_24_17,0,0,0,Theta_24_21,0,0,0,0,0;
    0,0,0,0,0,0,0,0,0,0,0,0,0,0,0,0,0,0,0,0,0,Theta_25_22,0,0,0,0;
    0,0,0,0,0,0,0,0,0,0,0,0,0,0,0,0,0,0,0,0,0,0,Theta_26_23,0,0,0;
    0,0,0,0,0,0,0,0,0,0,0,0,0,0,0,0,0,0,0,0,0,0,0,Theta_27_24,0,0;
    0,0,0,0,0,0,0,0,0,0,0,0,0,0,0,0,0,0,0,0,0,0,0,0,0,Theta_28_26;
    0,0,0,0,0,0,0,0,0,0,0,0,0,0,0,0,0,0,0,0,0,0,0,0,0,0;
    0,0,0,0,0,0,0,0,0,0,0,0,0,0,0,0,0,0,0,0,0,0,0,0,0,Theta_30_26];
    
    %Get Jacobian and eigenvalues

    J = LAMBDA * THETA;

    solution = eig(J);
    eigreal = real(solution);
    Parameters(run_n,:) = [F,P,D,A,Theta_1_13,Theta_2_1,Theta_3_2,Theta_4_3,Theta_5_3,Theta_5_4,Theta_6_5,Theta_7_3,Theta_7_6,Theta_8_4,Theta_8_7,Theta_9_8,Theta_10_3,Theta_10_9,Theta_11_11,Theta_12_10,Theta_13_12,Theta_14_3,Theta_15_6,Theta_17_14,Theta_18_15,Theta_19_16,Theta_20_17,Theta_21_18,Theta_22_19,Theta_23_20,Theta_24_17,Theta_24_21,Theta_25_22,Theta_26_23,Theta_27_24,Theta_1_25,Theta_28_26,Theta_30_26];
    result_eig(:,run_n) = solution;
    result_eigreal(:,run_n) = eigreal;
    
    
    %if rem(run_n,x) == 0
    %    disp(run_n)
    %end
    
end

result_max_eigreal = max(result_eigreal)';

histogram(result_max_eigreal,13)
count = length(nonzeros(result_max_eigreal(result_max_eigreal<0)))

% Save Files (un-comment)

%writematrix(result_max_eigreal,'Max_eig.csv')
%writematrix(Parameters,'Parameters.csv')
%writematrix(result_eig','Eigenvalues.csv')
```
