## Supplementary material for "The trade-off function of photorespiration in a changing environment": pseudocode BnB

```
FUNCTION pso_bnb(skm_model,discreet)
      FOR i IN 1: number_particles
            Initialize particle(i).Position
            particle(i).Position(discreet.Position) = discreet.Value
            particle(i).Cost = skm_model.cost_f(particle(i).Position)
      END FOR
      FOR it IN 1: max_iteration
            FOR i IN 1:number_particles
                  Update particle(i).Velocity
                  Update particle(i).Position
                  particle(i).Position(discreet.Position) =
                  discreet.Value
                  particle(i).Cost =
                  skm_model.cost_f(particle(i).Position)
                  IF particle(i).Cost > particle(i).Best.Cost
                        Update Particel(i).Best
                  END IF
                  IF particle(i).Best.Cost > global_best.Cost
                        Update global_best
                  END IF
            END FOR
            p_stable(it) = global_best.Cost/ skm_model.iterations
            IF p_stable(it) > 0.995 OR p_stable(it) - 2.58 *
            SQRT(p_stable(it) + (1 -
            p_stable(it))/skm_model.iterations) < p_stable(it - 20)
                  BREAK FOR
            END IF
      END FOR
END FUNCTION
```

```
n_var = INPUT("Number of Parameters to optimize")
discreet_parameter_values = [0, 1, -0.99, -0.1]
FOR i in 1:4
      discreet.Position = 1
      discreet.Value = discreet_parameter_values(i)
      Node(i).Position = 1
      Node(i).Value = discreet.Value;
      Node(i).Solution = pso_bnb(skm_model,discreet);
END FOR
WHILE Node IS NOT EMPTY
      I = MAX_POSITION(Node.Solution)
      NodeLevel = Node(I).Position + 1
      IF NodeLevel < n_var
            FOR i IN 1:4
                  discreet.Position = [1 : NodeLevel]
                  discreet.Value = [Node(I).Value Discreat(i)]
                  Node(i + length(Node)).Solution =
                  pso_bnb(skm_model,discreet)
            END FOR
            REMOVE Node(I)
      ELSE
            FOR i IN 1:4
                  discreet.Position = [1 : NodeLevel]
                  discreet.Value = [Node(I).Value Discreat(i)]
                  solved_node(i + length(SolvedNode)).Solution =
                  skm_model.cost_f(discreet)
            END FOR
            Node(I) = []
            Best_sol = MAX(solved_node.Solution)
            Solution = [Node.Solution]
            IF BestSol/2000 < 0.99
                  Prune = find(Solution/2000 < (BestSol/2000) - 2.58 *
                  sqrt((BestSol/2000) * (1 - (BestSol/2000)) / 2000))
            ELSE
```

```
            Prune = find(Solution/2000 < 0.995)
        END IF
        Node(Prune) = []
    END IF
END WHILE
```
